# Functional characterisation of rare variants in genes encoding the MAPK/ERK signalling pathway identified in long-lived Leiden Longevity Study participants

**DOI:** 10.1101/2023.06.01.541708

**Authors:** Helena Hinterding, Maarouf Baghdadi, Thies Gehrmann, Pasquale Putter, Mara Neuerburg, Nico Lakenberg, Erik B. van den Akker, P. Eline Slagboom, Joris Deelen, Linda Partridge

**Affiliations:** Max Planck Institute for Biology of Ageing, Cologne, Germany; Cologne Excellence Cluster on Cellular Stress Responses in Ageing-Associated Diseases (CECAD), University of Cologne, Cologne, Germany; Molecular Epidemiology, Department of Biomedical Data Sciences, Leiden University Medical Center, Leiden, The Netherlands; Leiden Computational Biology Center, Leiden, The Netherlands; Delft Bioinformatics Lab, Delft University of Technology, Delft, The Netherlands; Institute of Healthy Ageing, Department of Genetics, Evolution and Environment, University College London, London, UK

**Keywords:** MAPK/ERK signalling pathway, genetics, rare variants, longevity, ageing, functional characterisation, genome editing

## Abstract

Human longevity, which is coupled to a compression of age-related disease, has been shown to be heritable. However, the number of identified common genetic variants linked to this trait remains small. This may indicate that longevity is, at least to some extent, determined by rare genetic variants that are potentially family-specific. We therefore investigated whole-genome sequencing data of long-lived families from the Leiden Longevity Study for family-specific variants. We identified variants residing in genes involved in the mitogen-activated protein kinase (MAPK) cascade, a lifespan-associated and evolutionarily conserved pathway emerging from studies in model organisms. We subsequently generated mouse embryonic stem cells (mESCs) harbouring these variants and conducted *in vitro* functional characterisation. Two variants, located in *NF1* (Phe1112Leu) and *RAF1* (Asp633Tyr), reduce MAPK/extracellular signal-regulated kinase (ERK) signalling pathway activity in mESCs. At the proteomic and transcriptomic level, we observed prominent changes that were shared (e.g. up-regulation of the ribosome) and opposing between the variants (e.g. down-regulation of mTORC1 signalling in the RAF1^Asp633Tyr^ variant cell line versus up-regulation in the NF1^Phe1112Leu^ variant cell lines). These metabolic changes were accompanied by an opposing effect of the variants on proliferation. Moreover, the RAF1^Asp633Tyr^ variant improved resistance to replication stress, while this was not the case for the NF1^Phe1112Leu^ variant. In conclusion, we identified two rare genetic variants in long-lived families that influence MAPK/ERK signalling in a manner that has previously been linked to increased lifespan in model organisms. Interestingly, we also observe some opposing and diverging effects between the variants, which indicates that they may either have some pleiotropic effects that are not relevant to longevity or that they target slightly different mechanisms to elicit their effects in mESCs. Our findings suggest that mESCs offer a good starting point for *in vitro* characterisation of rare genetic variants linked to human longevity and can be used to assess which of these variants to take forward to *in vivo* studies in model organisms.

## Introduction

Human ageing is a complex trait influenced by many different factors, including genes, lifestyle, environmental circumstances, and access to healthcare ^1^. Mainly due to advances in targeting the environmental component (through nutrition, hygiene, and medicine), human lifespan has greatly increased in the past two centuries ^2^. However, there has thus far not been a similar increase in healthspan, i.e. the number of years lived before the occurrence of the first chronic disease ^3^. This is reflected in a growing period of loss of function and morbidity at the end of life, especially in women ^4^. Long-lived individuals, especially those from long-lived families, could offer mechanistic insights into how to improve late-life health, as they exhibit a compression of age-associated morbidity ^5,6^. The longevity trait, defined as the capacity to reach exceptionally old ages (e.g. >90 years), is transmitted across generations in families where many members show a lifelong excess survival up to extreme ages despite sources of extrinsic mortality, such as epidemics, famine, or conflicts in the past two centuries ^7–11^. This is exemplified by data from the Leiden Longevity Study (LLS) ^12^, which showed that individuals from long-lived families display beneficial immune and metabolic profiles in middle age ^13–15^. Moreover, the long-lived siblings of this two generational-cohort provide a robust sample of individuals with an enrichment of the heritable component of longevity ^16^. Therefore, we used genomic data from these families to identify genetic variants that may contribute to mechanisms of late-life health.

Methods for identifying genetic variants associated with human longevity have thus far mainly focused on common variants using a population-based approach. This approach, including a global genome-wide association study (GWAS) ^17^, revealed a handful of loci associated with longevity, of which only *APOE* and *FOXO3* were consistently identified across epidemiological and family-based studies of ancestrally diverse populations ^17–19^. Given the sparse findings from the GWAS approach, we instead focused on rare variants with potentially larger effects that are not tagged by genome-wide genotyping arrays ^20^. Unlike the hypothesis-free GWAS approach, which aims to identify common genetic variants across the genome associated with a trait in humans, the candidate-gene approach is applied to examine how a specific biological pathway relates to the trait under investigation ^21^. Using this approach, and under the assumption that some longevity genes may be evolutionary conserved, several groups have already started investigations in humans of lifespan-associated pathways and genes identified in model organisms ^22–26^.

The mitogen-activated protein kinases (MAPKs) are a family of highly conserved protein Ser/Thr kinases that integrate a wide variety of extracellular signals into a range of intracellular responses ^27^. There are multiple MAPK pathways in eukaryotic cells (e.g. up to seven in mammals), which coordinate cell proliferation, survival, apoptosis, mitosis, growth, migration, and metabolism ^28–30^. The wide range of different, and sometimes opposing, signals resulting from the different MAPK cascades in the same cells raises the question of how specificity is achieved. Many possible mechanisms have been proposed, such as manipulating duration or strength of signal, scaffold proteins, and alternative localisation ^31,32^, which are likely tissue- and context-specific. We specifically investigated one of the conventional MAPKs, namely the extracellular signal-regulated kinase 1/2 (ERK1/2) pathway, as both genetic and pharmacological inhibition of ERK1/2 phosphorylation have been linked to lifespan extension in yeast, worms, fruit flies, and mice ^33–37^. Conventional MAPK cascades are made of a sequence of evolutionary conserved kinases. For the ERK1/2 module these are MAPKKK (ARAF, BRAF, and RAF1 (CRAF)), MAPKK (MAP2K1 (MEK1)/MAP2K1 (MEK2)), and MAPK (MAPK3 (ERK1)/MAPK1 (ERK2)). This MAPK cascade is activated through receptor tyrosine kinases (RTKs) that transduce extracellular cues into the cell through activation of rat sarcoma virus (RAS) GTPases. These RAS GTPases function as central signalling hubs that initiate a downstream phosphorylation cascade through RAFs, MEK1/2, and ERK1/2 proteins to ultimately activate a variety of erythroblast transformation specific (ETS) transcription factors ^33^. The exact function of many of these ETS transcription factors in mammals is still unclear. The activation of the MAPK/ERK cascade achieves specificity through temporal, spatial, and tissue-specific regulation ^33,38^. Phosphorylation of ERK1/2 at the Thr and Tyr residues in the heavily conserved protein domain is essential for enzymatic activity and can be used as an indicator of MAPK/ERK signalling pathway activity ^39^. Interestingly, the MAPK/ERK signalling pathway also influences another major cellular growth and survival pathway, namely phosphoinositide 3-kinase (PI3K)-protein kinase B (AKT) signalling ^40,41^, that functions upstream of MAPK. The PI3K-AKT signalling pathway plays a crucial role in activating mammalian target of rapamycin complex 1 (mTORC1), primarily through the activation of AKT at Ser 473, which subsequently inhibits the tuberous sclerosis complex (TSC)1-TSC2 complex from inhibiting mTORC1 activity ^42–44^. Several studies using either pharmaceutical approaches (e.g. rapamycin) or genetic models (e.g. ribosomal protein S6 kinase 1 (S6K1) knockout in mice) have shown that the inhibition of mTORC1 leads to an extension of lifespan ^45–48^.

To study the role of genetic variation in the MAPK/ERK signalling pathway in modulating human longevity, we employed a heterologous systems approach to investigate the functional effects of rare protein-altering variants (with a minor allele frequency (MAF) below 0.01% in the general population) observed in the genome of long-lived individuals from the LLS ^49^. We measured their effects on MEK1/2 and ERK1/2 phosphorylation, protein (i.e. proteomics) and transcript (i.e. RT-PCR) abundance, and stress resistance in mouse embryonic stem cells (mESCs). Notably, we observed that two variants, located in *NF1* and *RAF1*, showed decreased MAPK/ERK signalling pathway activity (i.e. MEK1/2 and ERK1/2 phosphorylation) in mESCs. This was accompanied by prominent changes at the proteomic and transcriptomic level, both shared (e.g. up-regulation of the ribosome) and opposing between the variants (e.g. down-regulation of mTORC1 signalling in RAF1 variant mESCs versus up-regulation in NF1 variant mESCs). In addition, we found that the variant in *RAF1* showed reduced proliferation and improved resistance to replication stress *in vitro*, while the variant in *NF1* showed enhanced proliferation. Hence, although both the variants in *NF1* and *RAF1* showed functional effects, our findings highlight the complex regulation of factors downstream in the MAPK/ERK signalling pathway, thus suggesting different avenues by which this pathway can contribute to human longevity.

## Results and Discussion

### Identification of rare protein-altering variants in genes involved in the MAPK/ERK signalling pathway in long-lived families from the LLS

To investigate whether genetic variation in the MAPK/ERK signalling pathway plays a role in human longevity, we created a novel pipeline for the functional characterisation of rare variants detected in whole-genome sequencing (WGS) data of long-lived individuals from the LLS ^49^. The LLS cohort includes two generations of family members from 421 families selected based on long-lived siblings living in the Netherlands at the time of inclusion (men aged 89 years or above; women aged 91 years or above) ^12^. For the purpose of this study, we focused on the parental generation of the LLS and, more explicitly, the families who showed the strongest evidence for familial genetic enrichment of longevity, as previously defined by van den Berg and colleagues ^16^. This means that at least two siblings within the family belonged to the top 10% longest-lived of their birth cohort and had a parent meeting the same criterion ^16^. Of the 216 remaining families, we focused on the subset of 183 unrelated individuals from these families for whom WGS data was available. The mean age at death of these individuals is 98.2 years (standard deviation = 3.4 years). The genetic variants from the WGS data of the subset of individuals from long-lived families were subsequently filtered according to the following criteria (**Figure 1A**):

**(1)** The variant resides in the coding region of a MAPK/ERK signalling-related gene (see **Table EV1** for an overview of the gene list).
**(2)** The variant is protein-altering with a combined annotation-dependent depletion (CADD v1.7) score ≥20 (i.e. belonging to the top 1% of variants in the genome that are most likely to affect protein function) ^50^.
**(3)** The variant is absent (MAF = 0%) in Dutch-specific controls (BBMRI-NL) ^49,51^ and is absent or has a very low frequency (MAF <0.01%) in the general population, as determined using the publicly available reference database gnomAD (ALL population, https://gnomad.broadinstitute.org/ v.4.0.0).
**(4)** The variant is located in a genomic region that is shared between long-lived family members (identity-by-descent (IBD) ≥1 based on Illumina BeadChip data).
**(5)** The variant is validated using Sanger sequencing in the sequenced individual, as well as at least one long-lived sibling from the same family, to increase the likelihood that the variant contributes to familial longevity.

**Figure 1.**
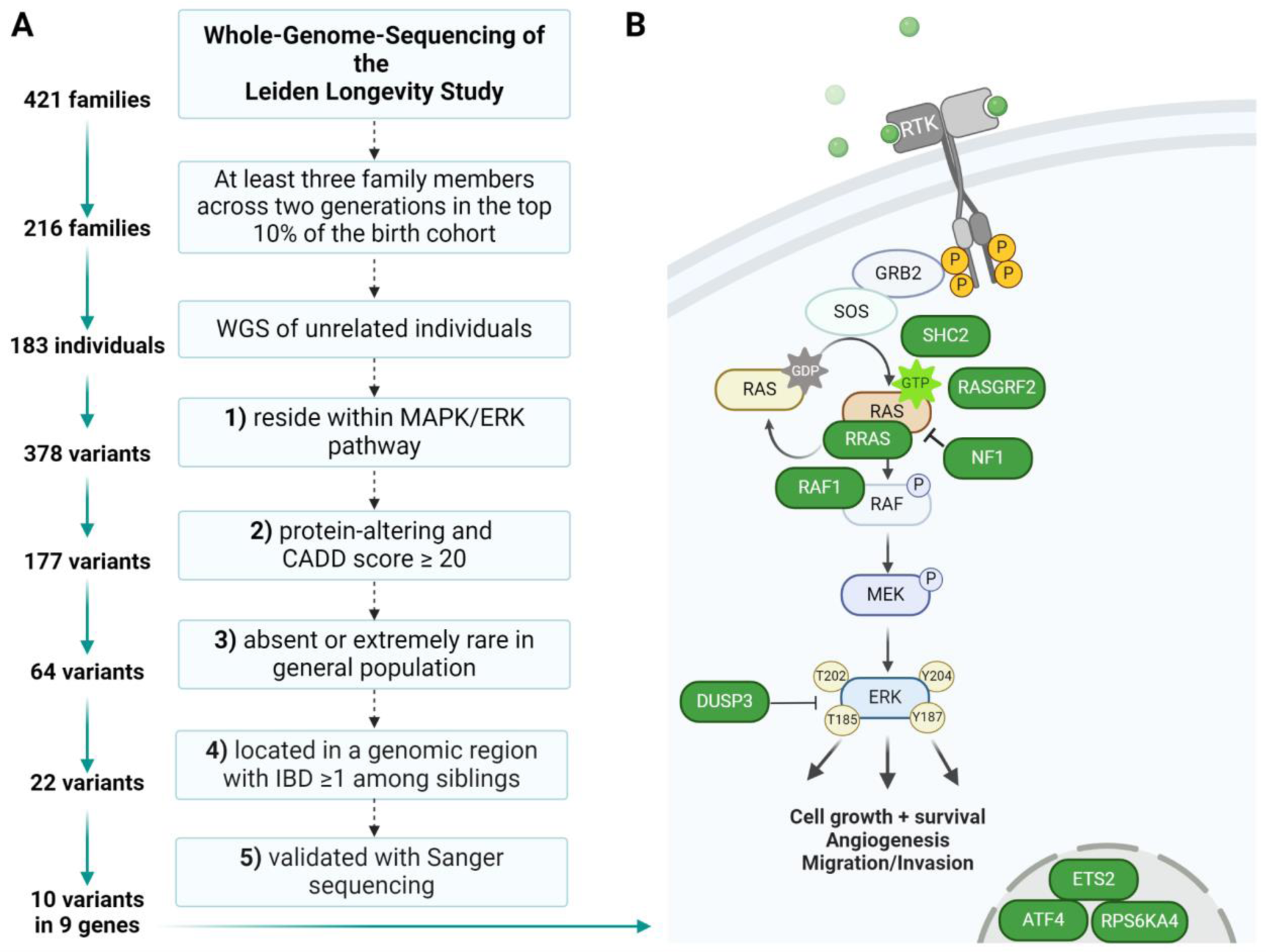
| Identification of MAPK/ERK signalling pathway-associated rare genetic variants in long-lived individuals. **(A)** Schematic of the pipeline used to filter the whole-genome-sequencing data of long-lived individuals from the Leiden Longevity Study. This filtering resulted in a list of 10 variants residing within 9 genes encoding the MAPK/ERK signalling pathway. **(B)** Simplified illustration of the MAPK/ERK signalling pathway with genes containing variants highlighted in green. The MAPK/ERK signalling pathway transmits cues into the cytoplasm and consists of upstream located RAS proteins and a phosphorylation cascade of RAF-, MEK-, and ERK-kinases. WGS; whole-genome sequencing, CADD; combined annotation-dependent depletion, IBD; identity-by-descent.

In total, we identified 10 variants meeting these criteria (**Figure 1B** and **Table 1**). All of these variants were present in the long-lived individuals in the heterozygous state. A limitation of our approach is that studies of rare variants are limited by statistical power ^52^. With this approach, a common issue faced by researchers is the inability to prove functionality and statistically significant enrichment in humans. We therefore only focused on variants that are shared among long-lived family members and have a very low frequency in the general population to increase the chance they actually contribute to the longevity of their carriers. Other groups have used similar approaches, but focused on variants identified in individuals without a known family history for longevity that were often also observed in control populations at a similar frequency ^23–25^. Although some of these variants show functional effects in cell lines ^23–25^, it is less likely that they contribute to longevity of their carriers, given that most of the controls will not reach an exceptionally old age. We therefore suggest that future functional studies of genetic variants should focus on genetic variants that are shared by long-lived individuals within families and are absent or have a very low frequency in the general population.

**Table 1.** | The 10 rare genetic variants in the MAPK/ERK signalling pathway identified in long-lived family members from the Leiden Longevity Study.

| Gene | Position | AA change in humans | AA change in mice | CADD score | Allele frequency in Europeans |
| --- | --- | --- | --- | --- | --- |
| *DUSP3* | 17-43769695-G-A | Arg158Cys | Arg158Cys | 32.0 | 0.00002203 |
| *RASGRF2* | 5-81073373-C-A | Leu270Ile | Leu270Ile | 26.9 | 0.000004496 |
| *ATF4* | 22-39522570-C-T | Arg342Cys | Arg340Cys | 27.3 | 0.00003981 |
| *SHC2* | 19-436409-T-C | Tyr266Cys | Tyr244Cys | 26.5 | 0.000009409 |
| ***RAF1*** | **3-12584564-C-A** | **Asp633Tyr** | **Asp633Tyr** | **27.1** | **0** |
| *ETS2* | 21-38818562-C-T | Arg243Trp | Arg95Trp | 23.2 | 0.000007627 |
| *RPS6KA4* | 11-64368241-C-T | Ala394Val | Ala394Val | 25.4 | 0.000001313 |
| ***NF1*** | **17-31232759-C-T** | **Ala1125Val** | **Ala1127Val** | **24.0** | **0.000008993** |
| ***NF1*** | **17-31232713-T-C** | **Phe1110Leu** | **Phe1112Leu** | **24.3** | **0.000002857** |
| ***RRAS*** | **19-49636720-C-T** | **Glu118Lys** | **Glu118Lys** | **33.0** | **0.00002373** |

Variants found in genes highlighted in bold were generated in mouse embryonic stem cells. CADD; combined annotation dependent depletion, AA; amino acid. The position of the variants is based on the human assembly GRCh38. The CADD scores are based on model GRCh38-v1.7. Allele frequencies are based on high quality genotypes of Europeans (non-Finnish) who are included in the gnomAD database v4.0.0.

### Generation of mESCs harbouring the identified variants

To study the functional consequences of the identified variants, we performed CRISPR/Cas9 gene editing using the Cas9^D10A^ nickase enzyme (double-nicking strategy) ^53^ to generate genetically engineered haploid AN3-12 mESCs ^54^. We opted for AN3-12 mESCs given that these cells have successfully been used for CRISPR/Cas9 genome editing in previous studies^55^. This mESC line provides a valuable tool for functional genomics, as it allows for precise genome editing in the haploid state and does not possess karyotypic abnormalities, such as polyploidy, that could affect the subsequent functional characterisation. Haploid mESCs can be edited more efficiently because targeted mutagenesis in diploid cell lines often results in unwanted insertions or deletions on at least one of the targeted alleles ^56^. Haploid mESCs naturally become diploid over time so, after the mutations had been introduced, we were able to select the diploid mESCs in which we studied the functional effects of the mutations in the homozygous state. However, a limitation of this approach is that the variants found in long-lived individuals were identified in the heterozygous state. As we expected mild functional effects, we aimed to increase their functionality by first generating homozygous cell lines that contain two alleles rather than one. Previous longevity interventions in the MAPK/ERK signalling pathway have focused on genes upstream of ERK1/2 ^34,36,37^. Therefore, we prioritised the variants located in genes upstream in the pathway and successfully generated mESC lines harbouring four of them, i.e. Ala1127Val and Phe1112Leu (for which we were able to generate two independent cell lines) in NF1, Asp633Tyr in RAF1, and Glu118Lys in RRAS (**Figure EV1**)^57–59^.

### MAPK/ERK signalling pathway activity of mutant mESC lines

Neurofibromin 1 (NF1), is a RAS GTPase activating protein that inhibits RAS activity, thereby reducing downstream ERK1/2 phosphorylation, after stimulation with growth factors such as serum, EGF, or PDGF ^60,61^. RAF proto-oncogene serine/threonine kinase 1 (RAF1), also called c-RAF, is a MAPKKK with direct contact to RAS and is activated through a multistep process involving the dimerization of the RAF domain which then activates the MAPKKs MEK1/2 by phosphorylating their activation segment. In turn, MEK1/2 phosphorylate the activation segment of the MAPKs ERK1/2, which then relocates to various subcellular compartments to induce context-specific responses ^62,63^. To assess the effect of the engineered cell lines harbouring the candidate mutations on the function of the MAPK/ERK module of MAPK signalling, we therefore measured the ratio of phosphorylated (Ser217/221 MEK1/2) over total MEK1/2 and phosphorylated (Thr202/Tyr204 ERK1 and Thr185/Tyr187 ERK2) over total ERK1/2 ^62^. The two independent mESC lines harbouring the NF1^Phe1112Leu^ variant as well as the cell line harbouring the RAF1^Asp633Tyr^ variant displayed a significant reduction in phosphorylation of both MEK1/2 and ERK1/2 (Figure 2A). The mESC lines harbouring the other variant located in the *NF1* gene, NF1^Ala1127Val^, as well as the variant in RRAS, RRAS^Glu118Lys^, showed no change in ERK1/2 phosphorylation (data not shown). We therefore decided to not take these variants forward for further experiments.

**Figure 2.**
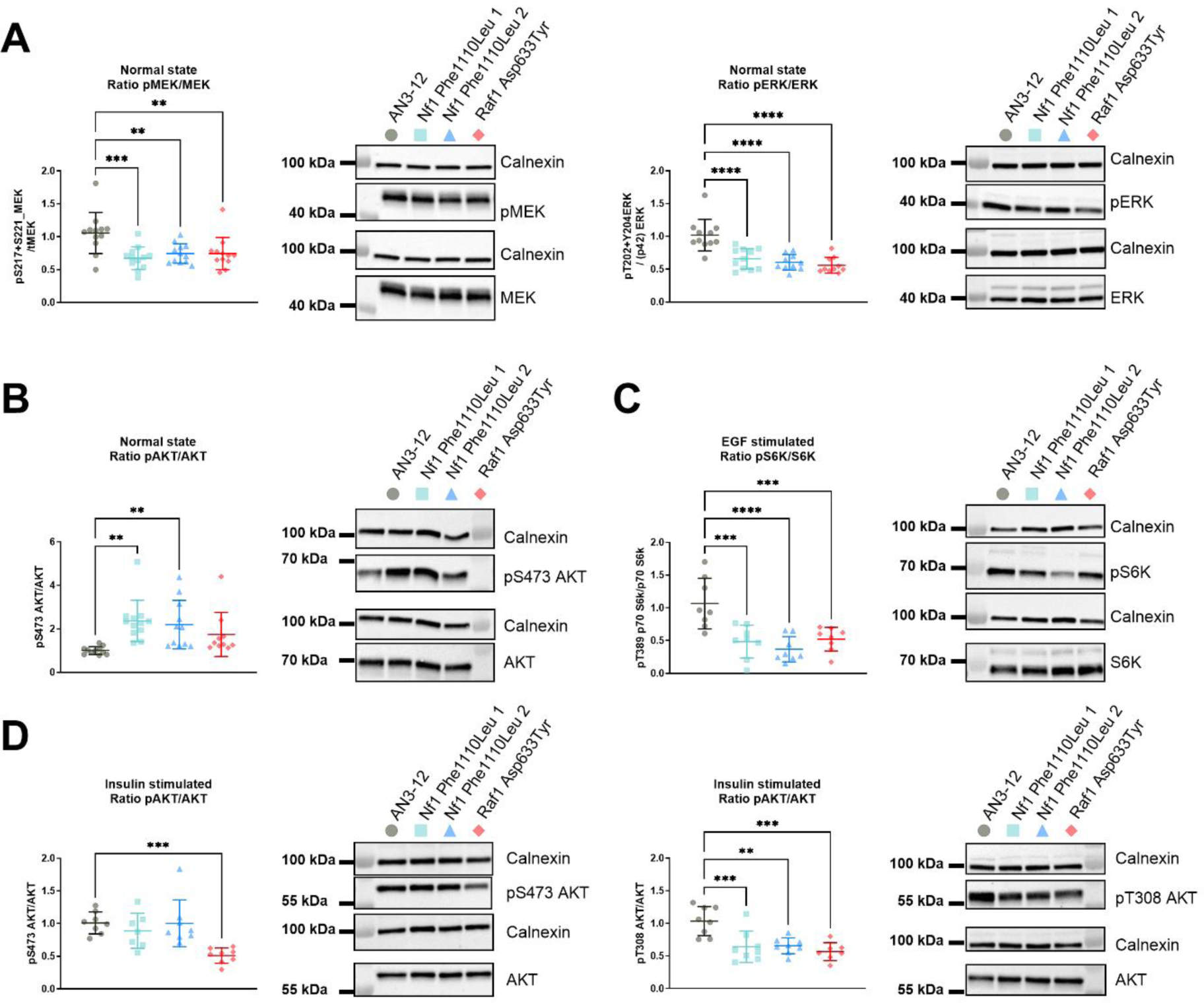
| Rare genetic variants in *NF1* and *RAF1* regulate MAPK/ERK and mTOR pathway activity in mESCs. **(A)** Significant reduction of phosphorylated MEK1/2 (Ser217/221) over total MEK1/2 and phosphorylated ERK1/2 (Thr202+Tyr204) over total ERK1/2 in the mESC lines harbouring the NF1^Phe1112Leu^ and RAF1^Asp633Tyr^ variants under normal growth conditions. (**B**) Significant reduction of phosphorylated AKT (Ser473) over total AKT in the mESC lines harbouring the NF1^Phe1112Leu^ variants under normal growth conditions. (**C**) Significant reduction of phosphorylated p70 S6K (Thr389) over total p70 S6K in the mESC lines harbouring the NF1^Phe1112Leu^ and RAF1^Asp633Tyr^ variants after EGF stimulation (100 ng/ml for 2 min.). (**D**) Significant reduction of phosphorylated AKT (Ser473 (left panel) and Thr308 (right panel)) over total AKT in the mESC lines harbouring the NF1^Phe1112Leu^ (Thr308 only) and RAF1^Asp633Tyr^ (both Ser473 and Thr308) variants after insulin stimulation (100 nM for 10 min.). For all experiments, Calnexin was used for normalisation. The data shown is from three (**A**) or two (**B-D**) independent experiments with four technical replicates each. Error bars represent standard deviation. Data was analysed using a one-way ANOVA and Dunnett’s post hoc test. *P < 0.05, **P < 0.01, ***P < 0.001, ****P < 0.0001.

### PI3K-AKT signalling pathway activity of generated mESC lines

In order to assess if the NF1^Phe1112Leu^ and the RAF1^Asp633Tyr^ mutations affected PI3K-AKT signalling pathway activity, we used the ratio of S6K phosphorylation (Thr389) over total S6K and phosphorylated AKT (Ser473) over total AKT as a readout for mTORC1 and mTORC2 activity, respectively ^64^. Unfortunately, we could not detect a reliable signal for S6K phosphorylation when mESCs were maintained in normal growth medium (normal state). However, both independent NF1^Phe1112Leu^ variant cell lines showed a significant elevation of phosphorylated AKT (Ser473) over total AKT, indicative of up-regulated PI3K-AKT and mTORC2 activity ^64^ (Figure 2B). This was unexpected, as an increase in NF1 activity is known to inhibit RAS function and thus result in a reduction of MEK1/2 and ERK1/2 phosphorylation, which should reduce PI3K recruitment and therefore reduce phosphorylated AKT (Ser473). However, previous studies have shown that mTORC1 inhibitors (such as rapamycin) can lead to increased phosphorylation of AKT (Ser473) due to the relief of negative feedback and up-regulation of upstream PI3K signalling ^65,66^.

### MAPK/ERK and PI3K-AKT signalling pathway activity after stimulation with EGF and insulin

In order to robustly assess any differential effects of our identified mutations on mTORC1/2 activity, we serum-starved the engineered mESC lines and subsequently stimulated them with EGF, PDGF, or insulin to acutely activate the MAPK/ERK and PI3K-AKT signalling pathways through a variety of RTKs. We found that acute stimulation with 100 ng/ml of EGF for 2 minutes ^67,68^ was sufficient to robustly activate the MAPK/ERK signalling pathway, while acute stimulation with 100 nM insulin for 10 minutes was able to activate the PI3K-AKT pathway ^69,70^ (**Figure EV2**). We therefore went ahead with these conditions. The subsequent stimulation of the genetically engineered mESC lines with EGF or insulin resulted in similar results for both MEK1/2 and ERK1/2 as for the non-stimulated cells (**Figure EV3A-B**).

A previous study showed that NF1 is a potential regulator of mTOR, as mTORC1 is constitutively active in NF1 deficient cells through up-regulation of TSC2 activity caused by an increased RAS and PI3K activation ^71^. Other studies suggest that the RAS-mediated ERK1/2 phosphorylation leads to dissociation of the TSC1-TSC2 complex thereby regulating mTORC1 ^42,72^. In addition, these studies found that pharmacological inhibition of ERK phosphorylation increased TSC1-TSC2 complex activity leading to increased mTORC1 inhibition and reduced phosphorylation of S6K (at Thr389). AKT phosphorylation also leads to inhibition of TSC2 and therefore increased phosphorylation of S6K ^42^. Hence, we decided to assess whether the PI3K-AKT pathway was differentially activated in our engineered mESC lines (after insulin stimulation) to determine whether the observed downstream effects could be delineated from MAPK/ERK signalling pathway activity. We measured the activity of mTORC1/2 by assessing the ratio of phosphorylated (Ser473 or Thr308 AKT) over total AKT and phosphorylated (Thr389) over total S6K. Both the NF1^Phe1112Leu^ and RAF1^Asp633Tyr^ variant mESC lines showed a down-regulation of phosphorylated over total S6K after both EGF (Figure 2C) and insulin stimulation (**Figure EV3B**). The engineered mESC lines hence showed a reduced mTORC1 activity, which has previously been associated with increased lifespan in model organisms ^73–75^. Moreover, the RAF1^Asp633Tyr^ variant mESC line showed a decrease in phosphorylated (Ser473 and Thr308) over total AKT after insulin stimulation, while the NF1^Phe1112Leu^ variant mESC lines only showed decreased phosphorylation at Thr308 (Figure 2D). There was no significant difference in phosphorylated AKT (Ser473) in EGF stimulated cell lines (data not shown).

Our data thus suggests that the consistent down-regulation of MAPK/ERK in our engineered mESC lines under different cellular states led to reduced mTORC1 activity, which we speculate could be regulated through increased TSC1-TSC2 complex activity. In addition, the differential effect of the mutations on AKT phosphorylation (at Ser473) after stimulation with insulin suggests that the RAF1^Asp633Tyr^ mutation likely reduced mTORC2 activity through reduced PI3K-AKT activity ^42^. Typically, inhibiting S6K phosphorylation (Thr389) due to reduced activity of mTORC1 results in up-regulation of phosphorylated AKT (Ser473) as mTORC1-dependent feedback is reduced. However, the simultaneous inhibition of AKT (Ser473) and S6K (Thr389) phosphorylation observed in the RAF1^Asp633Tyr^ variant cell line suggests that both mTORC1 and mTORC2 were inhibited. This is consistent with reports using AZD8055 an ATP-competitive inhibitor of mTOR kinase that inhibits both mTORC1 and mTORC2 ^76^. Moreover, the TSC1-TSC2 complex has no known effect on basal or insulin mediated phosphorylation of Akt (at Ser473 or Thr308) ^42^. Therefore, the significant down-regulation of phosphorylation of Akt (at Thr308) suggests that both the NF1^Phe1112Leu^ and RAF1^Asp633Tyr^ variants affected PI3K-AKT signalling upstream of TSC1-TSC2 (Figure 2D).

Taken together, the down-regulation of MEK1/2 and ERK1/2 phosphorylation combined with the downregulation of S6K phosphorylation suggests that NF1 function is increased in the NF1^Phe1112Leu^ variant cell lines (gain-of-function mutation). On the other hand, the data for the RAF1^Asp633Tyr^ variant cell line suggests a possible decrease in RAF1 activity (loss-of-function), given that human loss-of-function mutations in this gene result in reduced pathway activity ^77^, while gain-of-function mutations have shown the opposite effect ^78,79^.

### Proteomics of the NF1 and RAF1 mESC lines

The MAPK/ERK signalling pathway contains multiple positive and negative feedback loops, complicating the interpretation of functional readouts of their activity. We therefore also assessed broader effects of the mutants on the cellular proteome. We performed LC-MS/MS-based TMT-labelled quantitative proteomics of the mESC lines harbouring the NF1^Phe1112Leu^ and RAF1^Asp633Tyr^ variants. In total, we were able to detect 7726 proteins. Principal component analysis indicated that the two independent cell lines harbouring NF1^Phe1112Leu^ clustered together (**Figure EV4**). This decreased the likelihood that the observed effects in our genetically engineered mESC lines were caused by potential off-target effects inherent to genome editing. Moreover, the NF1^Phe1112Leu^ and RAF1^Asp633Tyr^ variants showed a distinct proteomic profile from each other and from the unedited (wildtype) cells (**Figure EV4**). The overlap in differentially abundant proteins (DAPs), i.e. those with a FDR-adjusted P<0.05, in the engineered mESC lines were strongest upstream in the MAPK/ERK signalling pathway (Figure 3A-B), characterised by a significantly increased abundance of EGFR and GRB2 and a decreased abundance of RASGRP4 and SHC1. The NF1^Phe1112Leu^ and RAF1^Asp633Tyr^ proteomic profiles started to diverge at the level of the RAS protein family, exemplified by an increased abundance of RRAS in the NF1^Phe1112Leu^ variant mESC line and decreased abundance in the RAF1^Asp633Tyr^ variant mESC line. The majority of the MAPK/ERK signalling pathway-specific DAPs in the RAF1^Asp633Tyr^ variant mESC line were down-regulated, suggesting negative feedback from ERK1 to upstream located proteins ^80^. Interestingly, RAF1 itself was also down-regulated in this mESC line, while NF1 was up-regulated, indicative of an overall reduced activity of the MAPK/ERK signalling pathway ^60–63,71^. On the other hand, NF1 expression was not affected in the NF1^Phe1112Leu^ variant mESC line.

**Figure 3.**
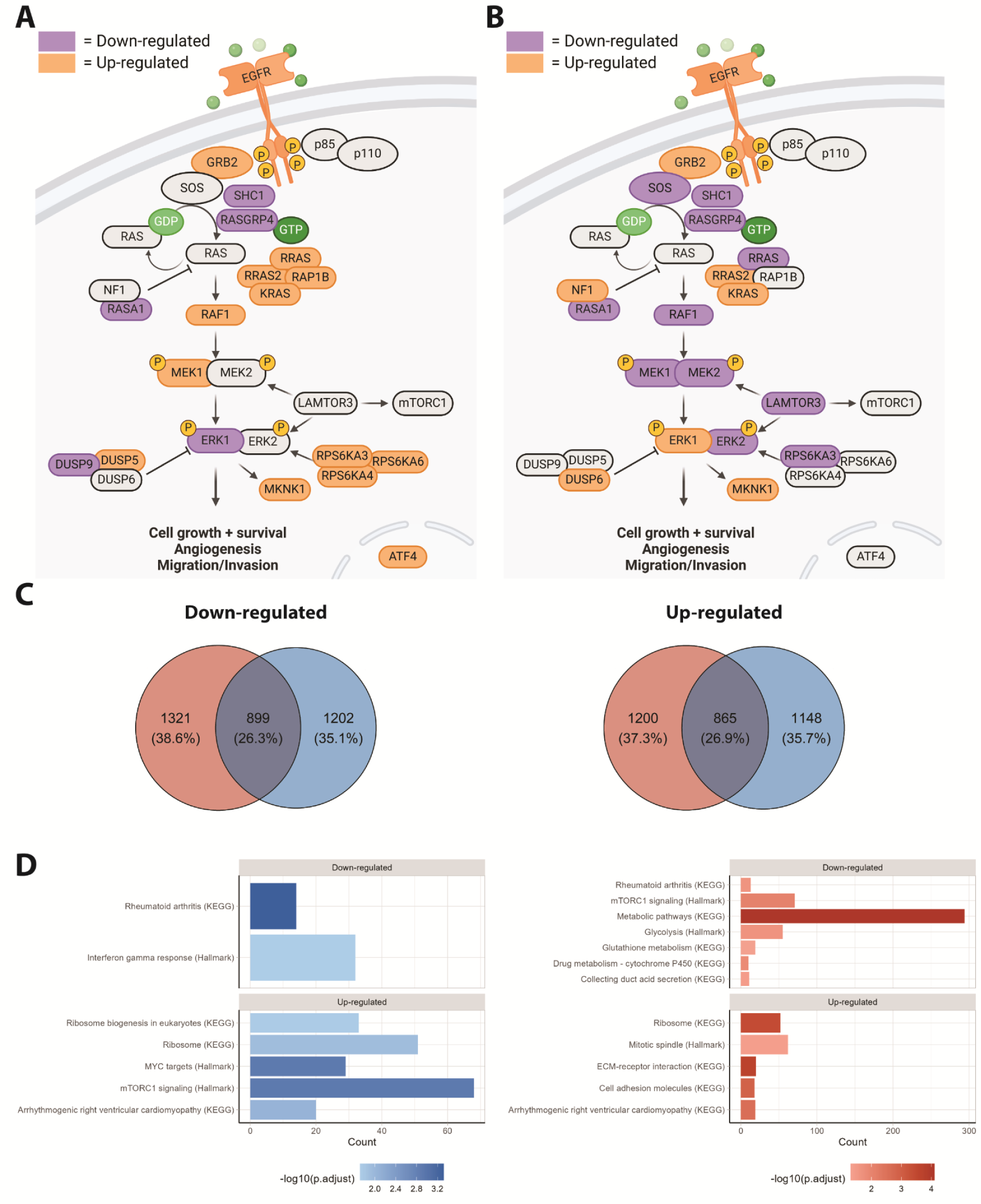
| Proteomics of the NF1^Phe1112Leu^ and RAF1^Asp633Tyr^ variant mESC lines. (**A-B**) Differentially abundant proteins (DAPs) in the MAPK/ERK signalling pathway in the NF1^Phe1112Leu^ (**A**) and RAF1^Asp633Tyr^ variant mESC line (**B**). (**C**) Venn diagrams representing the number and proportion of up- and down-regulated proteins in the NF1^Phe1112Leu^ and RAF1^Asp633Tyr^ variant mESC lines (FDR-adjusted P<0.05). (**D**) Overrepresentation analysis using KEGG and MSigDB pathways for DAPs (FDR-adjusted P<0.05) in NF1^Phe1112Leu^ and RAF1^Asp633Tyr^ variant mESC lines. The data is based on four technical replicates per cell line. Given the two independently generated NF1^Phe1112Leu^ variant mESC lines form one cluster in the PCA (**Figure EV4**), we decided to analyse them together (i.e. n = 8).

To probe effects of the variants on broader biological processes, we performed overrepresentation analysis using KEGG and Molecular Signatures Database (MSigDB) hallmark gene sets on the DAPs of the NF1^Phe1112Leu^ and RAF1^Asp633Tyr^ variant mESC lines (Figure 3C). For NF1^Phe1112Leu^, we observed a significant enrichment for rheumatoid arthritis and IFN-γ response in the down-regulated DAPS and for ribosome, mTORC1 signalling and MYC targets in the up-regulated DAPS (Figure 3D, **Table EV2**). For the RAF1^Asp633Tyr^ down-regulated DAPs, we detected strong enrichment of metabolic pathways and mTORC1 signalling, while the up-regulated DAPs show the strongest enrichment for ECM-receptor interaction and ribosome (Figure 3D, **Table EV2**). This again indicates that both mutants show shared, i.e. for rheumatoid arthritis and ribosome, but also opposing, i.e. for mTORC1 signalling, and differential effects at the proteome level.

### Transcriptional regulation of MAPK/ERK targets implicated in longevity in model organisms

We next performed RT-PCR on transcription factors previously linked to longevity in model organisms, i.e. Myc (*c-Myc*) ^81^, Foxo (*Foxo3*) ^82^, Nrf2 (*Nfe2l2*), and Ets transcription factors (*Ets1*, *Ets2*, *Etv1*, *Etv4*, *Etv5*, and *Etv6*) ^32^, to assess whether our identified mutations had similar effects. Interestingly, *c-Myc* was significantly up-regulated in the NF1^Phe1112Leu^ variants cell lines, but not the RAF1^Asp633Tyr^ variant cell line (Figure 4A). This is consistent with previous results highlighting the role of MYC in regulation of RNA polymerase transcription, ribosomal biogenesis, and cell growth ^83^. On the other hand, both the NF1^Phe1112Leu^ and RAF1^Asp633Tyr^ variant mESC lines showed an up-regulation of *Foxo3* (Figure 4B), which has previously been linked with healthy ageing and longevity ^84^. This result is consistent with the down-regulation of ERK1/2 phosphorylation, as ERK has previously been shown to interact with and regulate FOXO3 activity ^85^. Previous studies in worms suggest that up-regulation of the FOXO3 homolog DAF-16 through the PI3K-AKT signalling pathway also activates the Nrf2 homolog SKN-1, and that the transcriptional network of both of these transcription factors contribute to the longevity and stress resistance observed under inhibition of this pathway ^86,87^. However, we detected a significant down-regulation of *Nfe2l2* (Figure 4C), which has previously been associated with a decreased lifespan in both fruit flies and mice ^88,89^, although these effects may be tissue- and context-specific. The ETS transcription factor superfamily, which includes activators and repressors, offers a complex way of regulating cellular processes downstream of RTK signalling. This is due to their shared ETS DNA-binding domain, which recognize the same core DNA sequence ^33^. *Pointed* (*pnt*) and *anterior open (aop/yan)* are two ETS transcription factors that have previously been implicated in lifespan regulation in fruit flies through overlapping and dFOXO-independent effects ^32^. In fruit flies the activation of the MAPK pathway triggers a lengthy cascade ultimately leading to the activation of *pnt*, a transcriptional activator, and the inactivation of *aop/yan*, a transcriptional repressor. Sequence alignment of *pnt* reveals that *Ets1* and *Ets2* are the closest mammalian orthologues ^90^. Ets1 and Ets2 are both ETS transcriptional activators that ensure proper development of mammals in a complex temporal- and spatial-regulated manner ^91^. Interestingly, in response to the reduction of the MAPK/ERK pathway in the NF1^Phe1112Leu^ and RAF1^Asp633Tyr^ variant lines, we observed significant opposing effects on *Ets1* and *Ets2* expression within and across variant cell lines. Previous studies have linked a*op/yan* to lifespan extension through overlapping and distinct functions of dFOXO ^82^, while the same is true for ETV6/TEL and FOXO3 in relation to the mediation of hematopoietic stem cell survival ^92,93^, suggesting that the functional connection between these two transcription factors may be evolutionary conserved. Interestingly, we also observed an up-regulation of the transcriptional repressor *Etv6/Tel*, a mammalian orthologue of *aop/yan* ^94^, in the NF1^Phe1112Leu^ and RAF1^Asp633Tyr^ variant lines, although this effect was only significant in the NF1^Phe1112Leu^ variant lines. Etv6 is directly phosphorylated by ERK and deactivated downstream of Ras signalling, leading to the inability of Etv6 to bind to DNA and repress expression of downstream ETS targets ^95^. While we did not assess Etv6 phosphorylation, our data suggests that even though one of the ETS transcriptional activators is up-regulated in the NF1^Phe1112Leu^ (i.e. *Ets2*) and RAF1^Asp633Tyr^ (i.e. *Ets1*) variant lines, they both show an up-regulation of the ETS transcriptional repressor *Etv6*, potentially leading to the inhibition of ETS gene targets and down-regulation of the ETS pathway.

**Figure 4.**
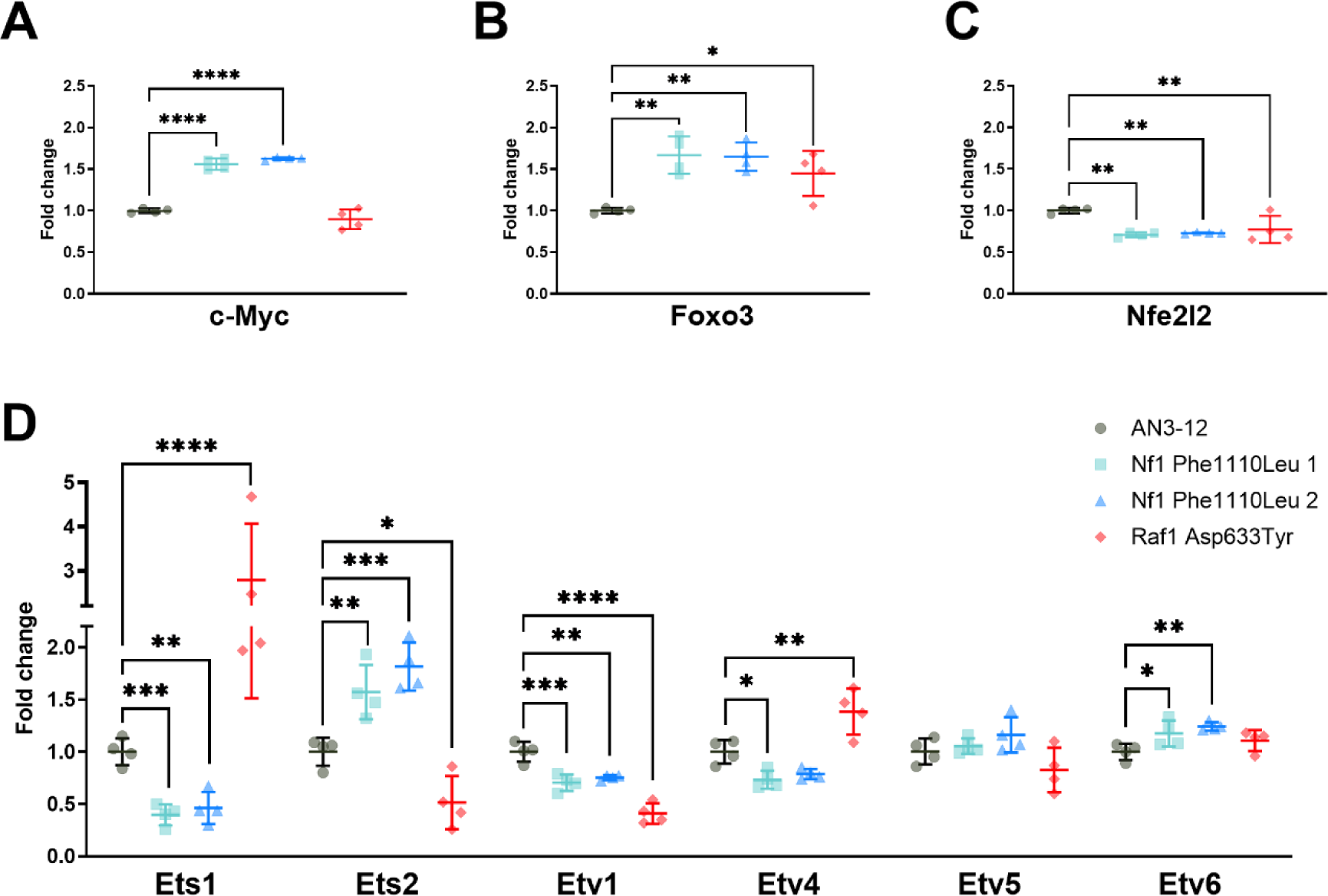
| Transcriptional regulation of MAPK/ERK targets implicated in longevity in model organisms. (**A-D**) Quantitative real-time PCR of *c-Myc* (**A**), *Foxo3* (**B**), *Nfe2l2* (**C**) and different Ets transcription factors (i.e. *Ets1*, *Ets2*, *Etv1*, *Etv4*, *Etv5*, and *Etv6*) in the NF1^Phe1112Leu^ and RAF1^Asp633Tyr^ variant mESC lines (**D**). Dots represent technical replicates (n = 4). Error bars represent standard deviation. Data was analysed using a one-way ANOVA and Dunnett’s post hoc test. *P < 0.05, **P < 0.01, ***P < 0.001, ****P < 0.001.

### Proliferation and resistance to replicative stress

Given the central role of the MAPK/ERK and PI3K-AKT signalling pathways in mediating cellular proliferation and growth, we decided to also asses proliferation of the genetically engineered cell lines ^43^. We observed that, in comparison to the wildtype mESCs, the proliferation of the NF1^Phe1112Leu^ variant mESC lines was increased, while that of the RAF1^Asp633Tyr^ variant mESC line was decreased under normal growth conditions (Figure 5A). The observed effects for NF1^Phe1112Leu^ variant cell line are in contrast with our western blot results, given that reduced MAPK/ERK signalling is normally associated with a reduction in growth. However, this discrepancy may be explained by the up-regulation of phosphorylated AKT (Ser473) in the NF1^Phe1112Leu^ variant mESC lines, which has been shown to mediate cell cycle progression and increase cell growth ^96^.

**Figure 5.**
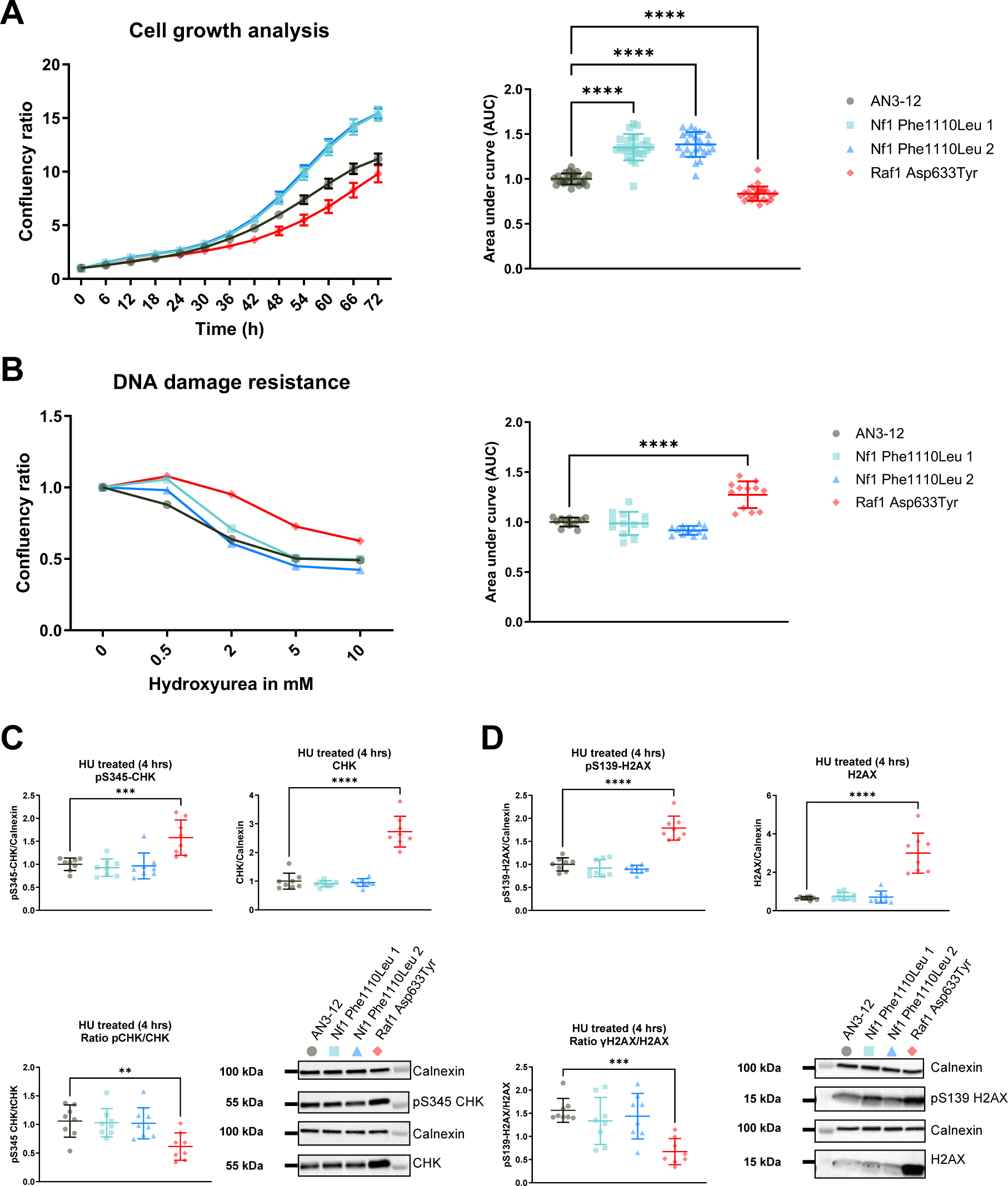
| Proliferation and resistance to replication stress of NF1^Phe1112Leu^ and RAF1^Asp633Tyr^ variant mESC lines. **(A)** The NF1^Phe1112Leu^ mESC lines show an increased proliferation, while the and RAF1^Asp633Tyr^ variant mESC line show the opposite under normal growth conditions. For each cell line, the confluency was normalised to the one observed at timepoint zero. The data is based on two independent experiments with 12 technical replicates each. The area under the curve (AUC) was calculated per experiment and normalised to that of the wildtype cells (right panel). **(B)** The RAF1^Asp633Tyr^ variant mESC line shows an increased resistance to replication stress after incubation with hydroxyurea for 22 hours. For each cell line, the confluency was normalised to the one observed for the untreated cells. The data shown is from three independent experiments with four technical replicates each. The area under the curve (AUC) was calculated per experiment and normalised to that of the wildtype cells (right panel). (**C**) Significant increase of phosphorylated CHK (Ser345) (left panel) and CHK (right panel) in the RAF1^Asp633Tyr^ mESC line after incubation with HU for 4 hours. (**D**) Significant increase of phosphorylated H2AX (Ser139) (left panel) and H2AX (right panel) in the RAF1^Asp633Tyr^ mESC line after incubation with HU for 4 hours. The data shown is from two independent experiments with four technical replicates each. Error bars represent standard deviation. Data was analysed using a one-way ANOVA and Dunnett’s post hoc test. *P < 0.05, **P < 0.01, ***P < 0.001, ****P < 0.001.

Previous studies have shown that the MAPK/ERK signalling pathway is activated in response to increased cellular stress to maintain survival ^97–100^. In order to assess if the NF1^Phe1112Leu^ and RAF1^Asp633Tyr^ variants also altered stress responses, we stressed the mESC lines harbouring them with hydroxyurea (HU). HU is an inhibitor of DNA synthesis and can lead to chromosomal damage (replication stress), such as double-strand breaks, depending on the concentration used, duration of treatment, and sensitivity of the cells. The two independent cell lines harbouring the NF1^Phe1112Leu^ variant did not differ from wildtype mESCs in their response to HU, while the RAF1^Asp633Tyr^ variant mESC line showed a significantly increased resistance (Figure 5B). Previous studies have shown that increased response to different stressors is often linked with increased lifespan and longevity ^101–104^, although this is not always observed ^105^. In order to assess how the variant cell lines regulated their response to HU, we looked at the effect on the DNA damage response (DDR) through western blotting. The DDR is a combination of multiple DNA pathways responsible for recognition, signalling, and repair of DNA damage in cells ^106^. CHK1 is a primary checkpoint kinase that regulates cell-cycle checkpoint signalling in response to DNA damage and replication blocks ^107^. To assess CHK1 activation after four hours of HU treatment, we quantified phosphorylated CHK1 (Ser345), the functional form of the kinase. Moreover, we assessed the levels of phosphorylated H2AX (Ser139), or γH2AX, which is required for DNA damage resistance through recruitment of DDR proteins to regions of double-strand breaks for repair ^108^. Consistent with the increased resistance to HU in the RAF1^Asp633Tyr^ variant mESC line (Figure 5B), we observed significantly higher levels of phosphorylated CHK1 (Ser345) (Figure 5C) and γH2AX (Figure 5D) indicating an increased recruitment of DDR chaperones, which may contribute to the observed replication stress resistance. We also observed a concomitant, but higher, increase in total CHK1 and H2AX protein levels after HU treatment (Figure 5C-D), which was in line with the proteomics data of the RAF1^Asp633Tyr^ variant mESC line measured under normal growth conditions. Taken together, this indicates that the observed resistance of the RAF1^Asp633Tyr^ variant mESC line to replication stress is partly mediated through increased activation of the DDR in response to HU and not only through decreased proliferation.

### Conclusion

In conclusion, we have created a pipeline for the identification and functional characterisation of rare genetic variants discovered in families enriched for longevity. We successfully applied this pipeline to determine the functional effects of genetic variation in the MAPK/ERK signalling pathway in haploid mESCs. This approach could also be applied to functionally characterise variants associated with other complex traits. We show that two of the identified variants, NF1^Phe1112Leu^ and RAF1^Asp633Tyr^, alter MAPK/ERK and PI3K-ERK signalling in mESCs in a manner that has previously been associated with increased lifespan in model organisms. However, we also observe some differential and opposing effects between the variants, which indicates that they lead to complex rewiring of the MAPK/ERK signalling pathway and potentially provide alternative strategies to elicit their effects in stem cells. Future studies should investigate the effect of the NF1^Phe1112Leu^ and RAF1^Asp633Tyr^ variants in the heterozygous state in both mouse and human cell lines, ideally obtained from carriers of the variants. Our results suggest that cellular models offer a good starting point for functional characterisation. However, in order to understand how MAPK/ERK signalling-related genetic variation may contribute to longevity, conserved variants should be studied in model organisms, such as mice, to be able to determine their more complex *in vivo* effects.

## Methods

### Study populations and selection criteria

The population used in this study was described previously ^12^. In brief, we utilised WGS data of 183 families belonging to the LLS. The LLS consists of Dutch families with at least two siblings meetings the following inclusion criteria: (1) men and women are at least 89 and 91 years old respectively, (2) participants have at least one living sibling who fulfils the first criterion, (3) the nonagenarian sibship share the same mother and father, and (4) the parents of the nonagenarian sibship are from European ancestry. For the purpose of this study, we applied a number of filters (see Figure 1A) to the WGS data of these nonagenarians, for which genetic variants were annotated with the Ensembl Variant Effect Predictor using the human assembly GRCh37 (hg19) as reference ^111^. To determine the frequency of the identified rare protein-altering variants in a background population similar to our long-lived individuals, we utilised sequencing data from 100 individuals of Dutch Caucasian origin (<=65 years old) collected by the Dutch Biobanking and Biomolecular Resources Research Infrastructure initiative (BBMRI-NL) ^51^. These individuals were not selected for particular characteristics other than that they should reflect a random sample of the apparently healthy Dutch population. The IBD between long-lived family members was calculated using the --ibd module from Merlin using previously generated genome-wide genetic data from Illumina BeadChips ^112,113^. The variants that remained after selection were remapped to human assembly GRCh38 (hg38) and the CADD scores and allele frequencies were updated based on newer versions of the used databases, i.e CADD GRCh38-v1.7 and gnomAD database v4.0.0, respectively.

### Culturing of mESCs

AN3-12 mESCs were generated previously ^54^ and were a kind gift from Martin Denzel. They were cultured in high glucose Dulbecco’s Modified Eagle Medium supplemented with 3.7 g/l Sodium bicarbonate, 13.7% FBS, 1% Pen-Strep, 1% L-Glutamate, 1% Sodium Pyruvate, 1% MEM Non-Essential amino acids, 0.1% beta-mercaptoethanol and 0.01% leukaemia inhibitory factor (mESC medium; Millipore, Massachusetts, United States). Cells were grown in petri dishes as semi-adherent colonies at 37°C with 5% CO2.

### Cloning

Guide RNAs (gRNAs) targeting the desired locus (**Table EV3**) were designed with Benchling (https://www.benchling.com) and ordered from Sigma-Aldrich (Missouri, United States). For cloning, Phusion High Fidelity Master Mix with GC Buffer (NEB, Massachusetts, United States) was used. All restriction digest reactions were performed with enzymes provided by NEB according to manufacturer’s instructions. T4 DNA Ligase (NEB) was used for ligation reactions. Two gRNAs were independently cloned into a vector containing the Cas9^D10A^ nickase enzyme (pSpCas9n(BB)-2A-GFP) (PX461), to minimise off target mutagenesis ^53^. pSpCas9n(BB)-2A-GFP (PX461) was a kind gift from Feng Zhang (Addgene plasmid #48140; https://www.addgene.org/48140/; RRID:Addgene_48140). Chemically competent OneShot TOP10 *Escherichia coli* (LifeTechnologies) were used for transformation of ligation reactions or plasmids according to manufacturer’s instructions. For positive selection of transformants, a 100 μg/mL ampicillin in Luria-Bertani (LB) plates were used. LB medium contained 5 g yeast extract, 10 g tryptone, 10 g NaCl 10 g (Sigma-Aldrich), and 1 l deionized water. The solution was adjusted to pH 7.0 and autoclaved before usage. For subsequent plasmid purification from bacteria, the QIAprep Miniprep or Midiprep Kits (Qiagen) were used. Cloning success was verified with Sanger Sequencing at Eurofins Genomics (Germany).

### CRISPR/Cas9 plasmid transfection

The created gRNA-containing vectors were subsequently transfected into the wildtype mESCs using Lipofectamine 3000 (Thermo Fisher Scientific, Massachusetts, United States) together with a short (120 bp) single-stranded DNA oligonucleotide (ssODN) (**Table EV4**). The ssODN contained the mutation of interest, as well as a minimum of two silent mutations. One silent mutation in the protospacer adjacent motif that prevents the Cas9n protein from re-cutting after successful repair as well as one mutation to add a restriction site for the screening of colonies.

### Fluorescence activated cell sorting (FACS)

We used the GFP that is co-expressed by the Cas9 plasmid (pSpCas9n(BB)-2A-GFP) (PX461) as a selection tool for transfected cells. Transfected mESCs were additionally stained with propidium iodide, to test for viability (Thermo Fisher Scientific), and Hoechst dye (Thermo Fisher Scientific), to assess chromosomal quantity, both according to manufacturer’s instructions. One day after transfection, haploid and GFP-positive cells were single-cell-sorted on 96-well plates using the BD FACSaria Fusion (BD Biosciences, New Jersey, USA) at the FACS and Imaging Core Facility at the Max Planck Institute for Biology of Ageing. Cells were grown for 10 days after which the single clones that revealed a positive edit after restriction digestion were genotyped.

### Genomic DNA preparation

Genomic DNA from mESCs was obtained by harvesting cells using 0.1% Trypsin (Thermo Fisher Scientific) and subsequent centrifugation at 500 rpm for 5 minutes for pellet formation. The cell pellet was resuspended in QuickExtract DNA Extraction Solution (Lucigen, United Kingdom) according to manufacturer’s instructions. Genomic DNA was stored at 4°C for downstream applications or at −20°C until further usage.

### Genotyping

For standard genotyping of the mESCs, GoTaq Master Mix (Promega, Wisconsin, United States) was used according to manufacturer’s instructions. Oligonucleotides (**Table EV5**) were ordered from Sigma-Aldrich. To analyse the size of DNA fragments or plasmids, 1x or 2x TAE (Tris base, acetic acid, ethylenediaminetetraacetic acid; Promega) buffered agarose gel electrophoresis was used to separate the DNA. DNA was stained with 10 μL Gel Red Stain (Invitrogen, Massachusetts, United States) per 100 mL agarose gel. Electrophoresis was performed with the Sub Cell GT horizontal electrophoresis system (Bio-Rad, California, United States) at 60-130 V for 30-60 minutes. Hyperladder 50 bp and 1 kb (both from Bioline, Tennessee, United States) were used as DNA fragment size markers.

### Stimulation with insulin, PDGF, and EGF

Cells were washed twice with warm PBS to remove any residual serum, after which they were serum-starved for 16 hours in mESC medium without FBS. This step reduces basal signalling activity and ensures that the observed effects are due to the stimulation. After serum starvation, cells were washed once with phosphate-buffered saline (PBS), after which the stimulant-containing starvation medium was added. Stimulation was performed by adding insulin (100 nM), PDGF (50 ng/ml or 100ng/ml) (520-BB, RNDSystems, Minnesota, USA), or EGF (25 ng/ml or 100ng/ml) (E4127, Merck, Germany) to cells for 10, 30, and 2 minutes, respectively.

### Protein extraction and western blotting

Before protein isolation, cells were washed twice with cold PBS (Sigma-Aldrich) after which 180-200 μl RIPA buffer (10 ml RIPA Buffer (Thermo Fisher Scientific), one cOMPLETE EDTA (Roche, Switzerland), and one PhosSTOP tablet (Roche)) was added to the samples on ice. Cells were harvested using a cell scraper after incubation them for 10 minutes. The suspension was sonicated for 10 seconds with 20% power for 10 cycles (one second on and one second off) pulses and centrifuged at 13000 rpm at 4°C for 15 minutes. Supernatant was transferred to a 1.5 ml tube and stored at −80°C until usage. The protein concentration was determined using the BCA Protein Assay Kit (Thermo Fisher Scientific) and samples were diluted accordingly. Laemmli Buffer (100 mM Tris pH 6.8, 20% glycerol, and 4% SDS (Carl Roth, Germany)) and 5% β-mercaptoethanol (Sigma-Aldrich) were added to the samples after which they were heated to 95°C for 10 minutes.

Equal amounts of proteins (20 μg) were loaded and separated at 100-150 V for 120 minutes, using Any kD™ Criterion™ TGX Stain-Free™ Protein Gel (Bio-Rad #5678124). Gels were activated with UV light for 2.5 minutes with the ChemiDocImager (Bio-Rad) before transferring them onto a 0.45 μm nitrocellulose membrane (GE Healthcare, Illinois, United States) at 100 V for 50 minutes. After the transfer, Stain-Free images were captured using the ChemiDocImager (Bio-Rad). The membrane was blocked using 5% non-fat dry milk powder in TBST for one hour and then incubated with primary antibodies (**Table EV6**) at 4°C overnight. Afterwards, the membranes were washed three times for 10 minutes each with TBST and the appropriate Horseradish peroxidase (HRP) coupled secondary antibody (1:10,000) was added (Thermo Fisher Scientific) in TBST containing 5% non-fat milk and incubated for 1 hour. The membranes were subsequently washed three times for 10 minutes each with TBST after which signal development was performed with ECL Select Western Blotting Detection Reagent (GE Healthcare). Images were acquired using the ChemiDocImager (Bio-Rad) and analysed with the Image Lab Software (Biorad).

### RNA isolation, cDNA synthesis, and RT-PCR

Total RNA was extracted from mESCs using TRIzol Reagent (Invitrogen, California, USA) according to the manufacturer’s instructions with slight modifications to optimize yield and purity. Briefly, samples were homogenised by pipetting in 0.4 mL of TRIzol reagent per well of mESCs. After homogenisation, samples were incubated at room temperature for 10 minutes. Chloroform (80 uL) was added, and tubes were shaken vigorously for 15 seconds and then allowed to stand at room temperature for 10 minutes. The mixture was centrifuged at 12,000x g for 15 minutes at 4°C. RNA was precipitated by mixing with isopropyl alcohol (0.2 mL) as well as 1 uL of GlycoBlue Coprecipitant (AM9515, Thermo Fisher Scientific) and samples were incubated at −80°C overnight. Samples were allowed to thaw then centrifuged at 12,000x g at 4°C for 10 minutes. The RNA pellet was washed once with 75% ethanol, vortexed, and centrifuged at 7,500x g at 4°C for 5 minutes. The RNA pellet was air-dried for 10 minutes and then dissolved in 20 uL RNase-free water. RNA was treated with deoxyribonuclease using DNA-free™ DNA Removal Kit (AM1906, Thermo Fisher Scientific) according to manufacturer’s instructions. The quantity and quality of RNA were assessed using a NanoDrop spectrophotometer (Thermo Fisher Scientific). Complementary DNA (cDNA) was prepared with the SuperScript III First-Strand Synthesis SuperMix (18080400, Thermo Fisher Scientific) from 4000 ng of RNA and diluted 1:5 for qRT-PCR. Samples consisting of cDNA mixed with the TaqMan Fast Advanced Master Mix (Thermo Fisher Scientific, 4444557) and TaqMan Assay probes were loaded in technical quadruplicates for qRT-PCR on a QuantStudio 6 Flex Real-Time PCR System (Thermo Fisher Scientific, 4485691). The ΔΔCt method was used to provide gene expression values after normalizing to the known reference gene *Gapdh*. Additional information regarding the used TaqMan qPCR Assays is provided in **Table EV7**.

### Proteomics

#### Protein extraction

mESCs were grown at 500,000, pellets collected by trypsinization and washed three times with PBS (Sigma-Aldrich). The cell pellet was resuspended in 20 μl lysis buffer (6 M guanidinium chloride, 2.5 mM Tris-HCL, 0.25 M Tris(2-carboxyethyl) phosphine and 0.8M chloroacetamide dissolved in purified MilliQ water (Merck)). The mixture was heated to 95°C for 10 minutes and sonicated in a Bioruptor (Diagenode, Belgium) (30 seconds sonication, 30 seconds breaks, 10 cycles). The lysate was centrifuged at 20,000x g for 10 minutes and transferred to new tubes. Protein concentrations were measured with NanoDrop (Thermo Fisher Scientific) and 300 μg diluted ten times with 20 mM Tris and digested with trypsin 1:200 overnight. The digestion was stopped the next morning by adding 100% formic acid (FA) and samples cleared by centrifugation at 20,000x g for 10 minutes.

#### Peptide cleaning

Peptide cleaning was performed with house-made 30 μg C18-SD StageTips. In brief, these tips were wet with methanol, then 40% acetonitrile/0.1% FA and equilibrated with 0.1% FA by table centrifugations of 1-2 minutes without letting the tips dry in between these steps. The total protein digest was then loaded in 0.1% FA and washed with 0.1% FA. Final elution was done with 100 μl 40% acetonitrile/0.1% FA at 300x g for 4 minutes and elutes dried in a SpeedVac at 45°C for 35 minutes. The digest was resuspended in 20 μl 0.1% FA and concentrations measured with NanoDrop (Thermo Fisher Scientific). 4 μg of peptides were used for subsequent TMT labelling.

#### Tandem Mass Tag (TMT) Pro labelling and fractionation

The peptides were reconstituted in 0.1 M triethylammonium bicarbonate buffer (Sigma Aldrich) and TMT Pro labelling done according to manufacturer’s instructions (Thermo Fisher Scientific). The ratio of peptides to TMT Pro reagent was 1:20. Labelled peptides were pooled, dried and resuspended in 200 μl 0.1% FA. These were then fractionated on a 1 mm x 150 mm ACQUITY column, packed with 130 Å, 1.7 µm C18 particles (Waters, Massachusetts, United States), using an Ultimate 3000 UHPLC (Thermo Fisher Scientific). Peptides were separated at a flow of 30 µL/min with an 88 min segmented gradient from 1% to 50% buffer B for 85 minutes and from 50% to 95% buffer A for 3 minutes. Buffer A contained 5% acetonitrile, 10 mM ammonium bicarbonate, buffer B contained 80% acetonitrile, 10mM ammonium bicarbonate. Fractions were collected every three minutes, pooled and dried in a SpeedVac.

#### LC-MS/MS analysis

Dried fractions were resuspended in 0.1% FA and separated on a 50 cm, 75 µm Acclaim PepMap column (Thermo Fisher) and analysed on a Orbitrap Lumos Tribrid mass spectrometer (Thermo Fisher) equipped with a FAIMS device (Thermo Fisher). The FAIMS device was operated in two compensation voltages, −50 V and −70 V. Peptide separations were performed on an EASY-nLC1200 using a 90 min linear gradient from 6% to 31% buffer. Buffer A contained 0.1% FA, buffer B contained 0.1% FA and 80% acetonitrile. The analytical column was operated at 50°C. Raw files were split based on the FAIMS compensation voltage using FreeStyle (Thermo Fisher). The final data was analysed using MaxQuant ^114^.

### Proliferation and replicative stress

The Incucyte® Live-Cell Analysis System was used to measure cell growth over time. To this end, 2,500 cells/well (96 well plate) were plated in 12 technical replicates per cell line. After allowing the cells to settle for 2 hours, plates were transferred into the Incucyte® Live-Cell Imager. Followed by a 3-hour waiting time, images were taken every 6 hours for 72 hours at a magnification of 10X using the phase image channel. Three images were taken per well. The AI tool from the Incucyte® Live-Cell Analysis System was used to determine the confluence. The average phase object confluence per well at any timepoint was normalised to the average phase object confluence per cell line at the first time point. For the assessment of replicative stress, 10,000 cells/well (96 well plate), with 16 replicates per cell line, were plated. Hydroxyurea (HU) was dissolved freshly at a concentration of 50 mg/mL in water and further diluted in serum free medium to the working stock concentrations. After letting cells settle for 3 hours, HU was added at the respective concentrations to 4 wells per cell line. The plates were moved into the Incucyte® Live-Cell Imager and 3 phase image channel pictures at a magnification of 10X per well were taken after 22 hours. The AI tool from the Incucyte® Live-Cell Analysis System was used to determine the confluence. The average phase object confluence of the three images per well was normalised to the average phase object confluence per cell line of the control without HU.

### Statistics

Data were analysed statistically using GraphPad Prism 9.5.0. The value of α was 0.05, and data were expressed as *P < 0.05; **P < 0.01; ***P < 0.001; ****P < 0.0001. Across group comparisons were made using a one-way ANOVA and between groups using a Dunnett’s post hoc test. R 4.2.2 was used for the proteomic analyses. The proteomic data was processed and analysed using a custom-made pipeline based on the R packages “edgeR”, “VennDiagram”, “clusterProfiler” (over-representation analyses), “msigdbr”, and “ggplot2”. All error bars correspond to standard deviations.

## Data availability

The code used for the analyses of the proteomics data is available upon request from the corresponding author Joris Deelen. The proteomics data produced in this study is available in the PRoteomics IDEntifications Database (http://www.ebi.ac.uk/pride/archive/projects/PXDXXXXXX).

## Acknowledgments

We thank the study participants of the Leiden Longevity Study. We thank the Proteomics and FACS and Imaging Core Facilities of the Max Planck Institute for Biology of Ageing for outstanding technical help and advice. We thank Isabell Brusius for assistance with the proteomics analyses. Figure 1 and Figure 3A-B were created with BioRender.com. This project has received funding from the European Research Council (ERC) under the European Union’s Horizon 2020 (grant agreement n° 741989) and Horizon Europe (ElucidAge, 101041331) research and innovation programmes. Views and opinions expressed are however those of the author(s) only and do not necessarily reflect those of the European Union or the European Research Council Executive Agency. Neither the European Union nor the granting authority can be held responsible for them.

## Author contributions

**Helena Hinterding:** Methodology; investigation; formal analysis; visualisation; writing – original draft; writing – review and editing.

**Maarouf Baghdadi:** Conceptualization; methodology; investigation; formal analysis; visualisation; writing – original draft; writing – review and editing.

**Thies Gehrmann:** Investigation; formal analysis.

**Pasquale Putter:** Investigation; formal analysis; validation; writing – review and editing.

**Mara Neuerburg:** Investigation; formal analysis.

**Nico Lakenberg:** Investigation.

**Erik B. van den Akker:** Supervision; methodology.

**P. Eline Slagboom:** Conceptualization; resources; supervision; funding acquisition; writing – review and editing.

**Joris Deelen:** Conceptualization; resources; supervision; methodology; investigation; formal analysis; funding acquisition; writing – original draft; writing – review and editing.

**Linda Partridge:** Conceptualization; resources; supervision; funding acquisition; writing – review and editing.

