## Supplementary Figures and Tables for "Functional characterisation of rare variants in genes encoding the MAPK/ERK signalling pathway identified in long-lived Leiden Longevity Study participants"

Hinterding H., Baghdadi M., et al.

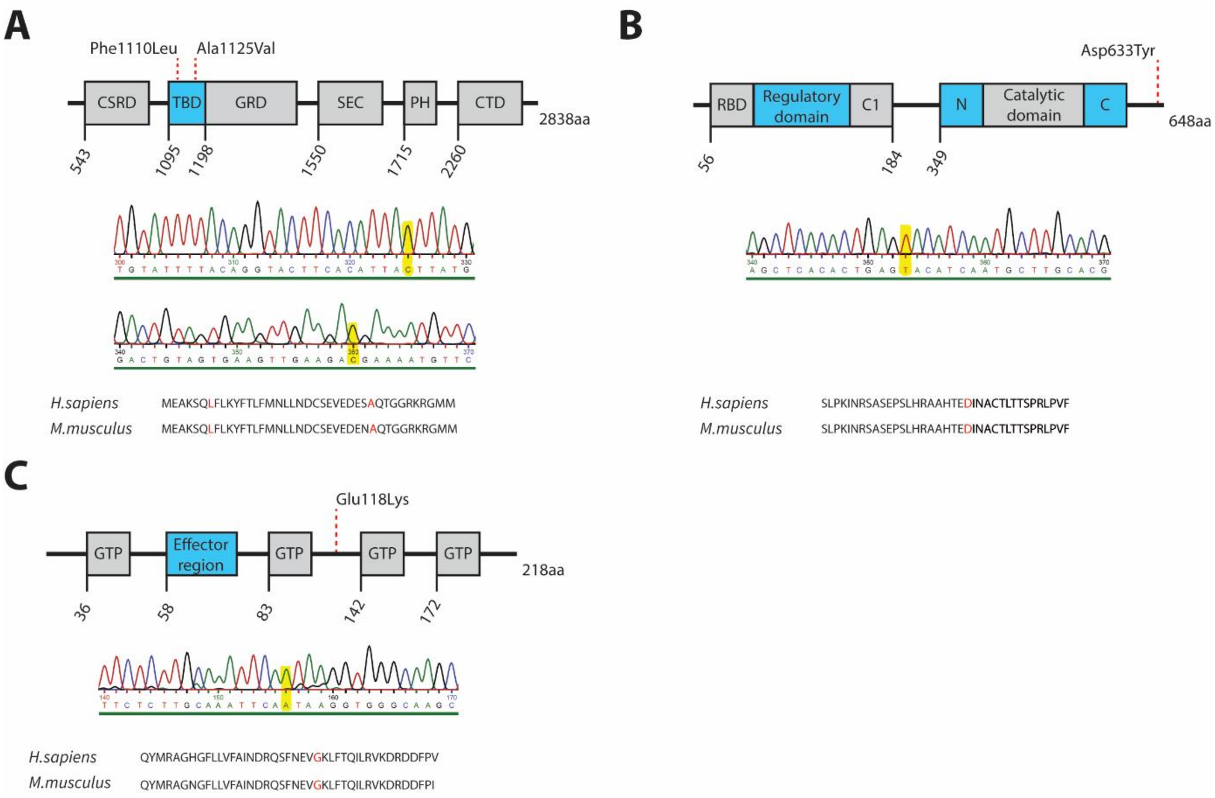


**Figure EV1 | Location and conservation of rare genetic variants in genes involved in the MAPK/ERK signalling pathway for which transgenic mESC lines were created.**

(**A-C**) Schematic representation highlighting the features of the NF1 (**A**), RAF1 (**B**), and RRAS (**C**) proteins, including the location of the identified variants (denoted by dashed red lines), the DNA sequence resulting from Sanger sequencing validation (with the mutation highlighted in yellow), as well as the sequence homology between humans and mice.

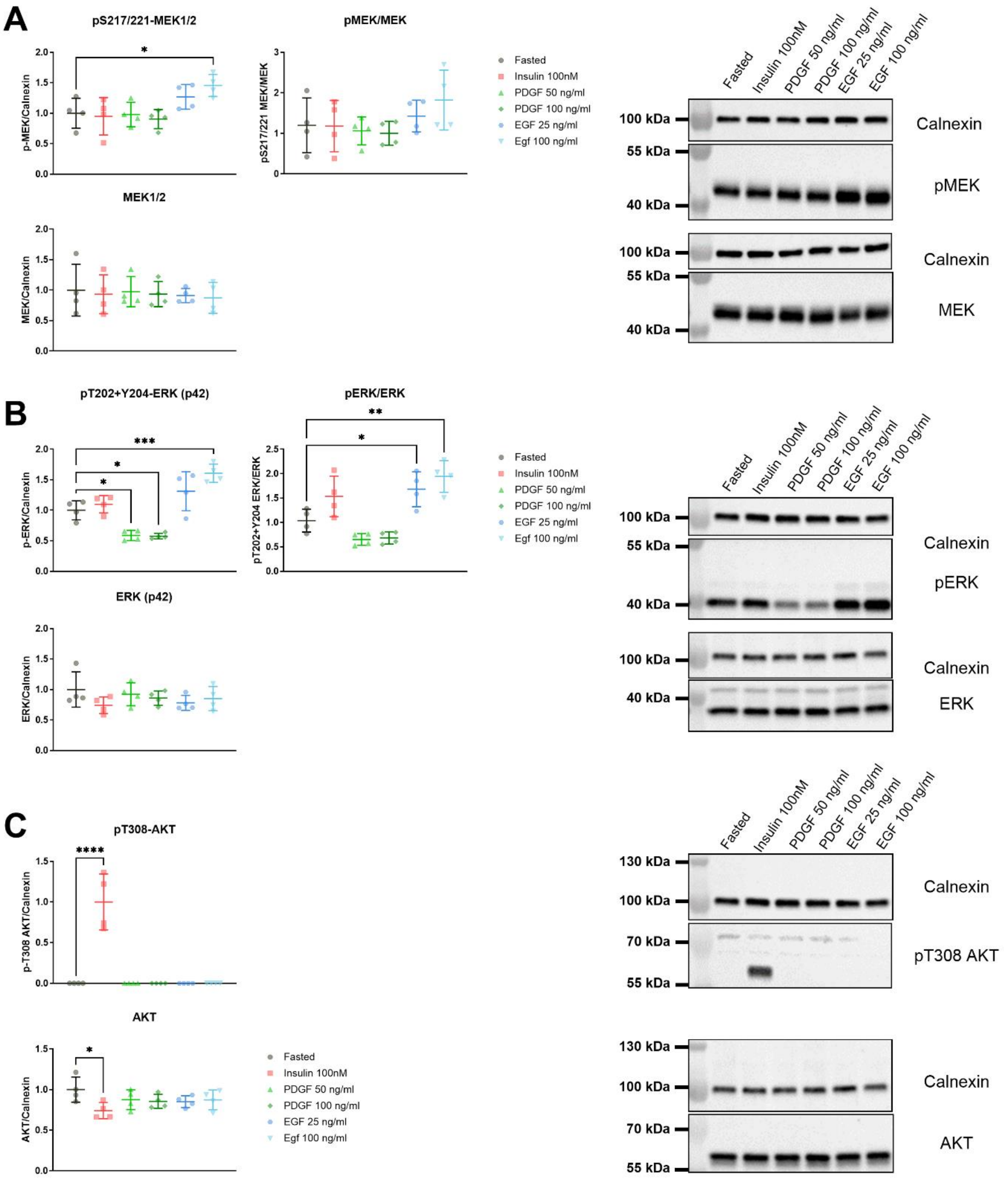


**Figure EV2 | MAPK/ERK and IIS pathway-related effects after insulin, PDGF, or EGF stimulation in wildtype mESCs.**

(**A**) EGF stimulation (100 ng/ml for 2 min.) was sufficient to significantly up-regulate phosphorylated MEK1/2 (Ser 217/221) over total MEK1/2 after fasting (6 hr.) in wildtype mESCs. (**B**) EGF stimulation (25 ng/ml and 100 ng/ml for 2 min.) was sufficient to significantly up-regulate the ratio of phosphorylated (T202+Y204) ERK1/2 over total ERK1/2 after fasting (6 hr.) in wildtype mESCs. (**C**) Insulin stimulation (100 nM for 10 min.) was the only growth factor to significantly up-regulate the phosphorylation of AKT (T308) over total AKT. Error bars represent standard deviation. Data was analysed using a one-way ANOVA and Dunnett's post hoc test. *P < 0.05, **P < 0.01, ***P < 0.001, ****P < 0.0001.

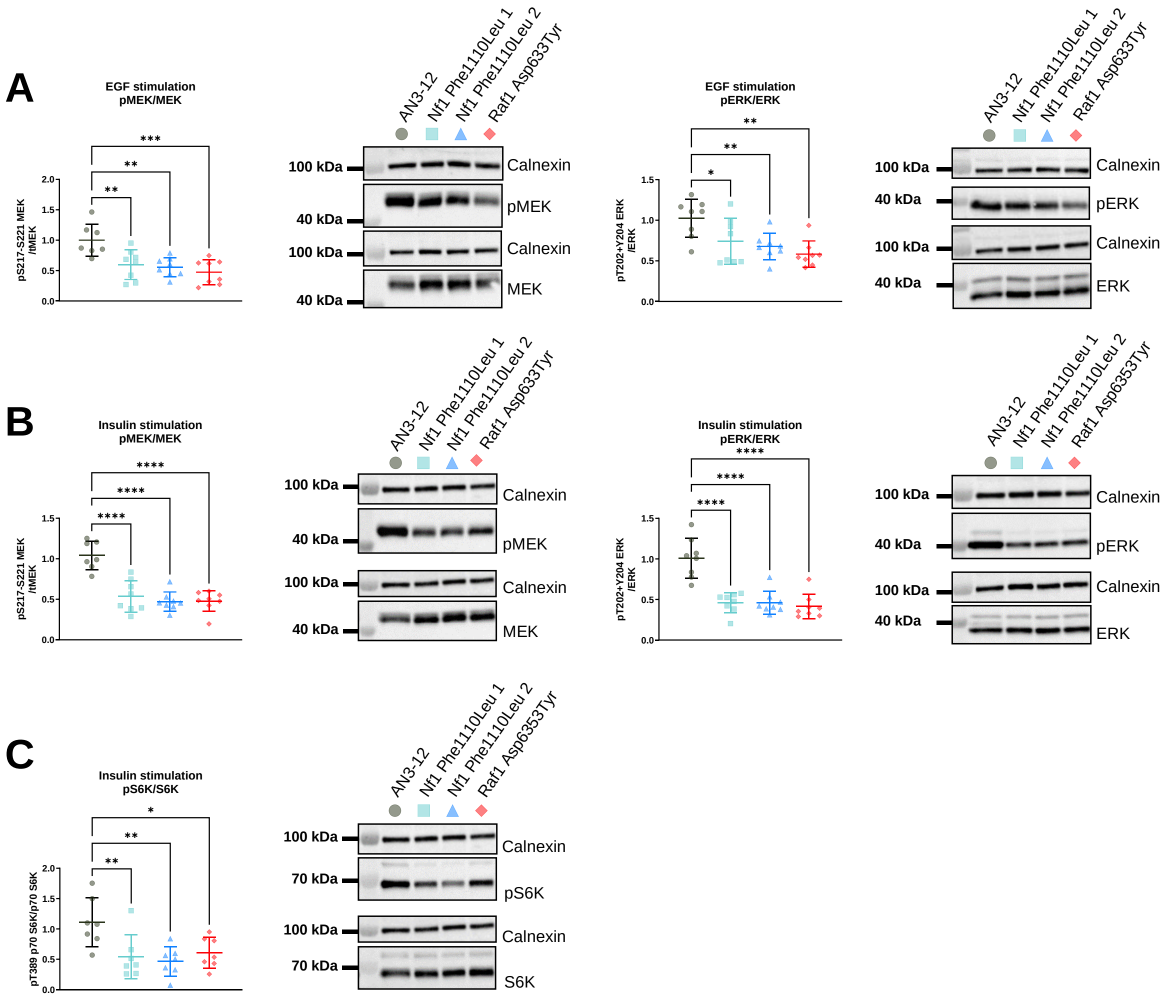


**Figure EV3 | MAPK/ERK and IIS pathway-related effects of the NF1$^{Phe1112Leu}$ and RAF1$^{Asp633Tyr}$ variant mESC lines after EGF and insulin stimulation.**

(**A**) Significant reduction of phosphorylated MEK1/2 (Ser217/221) over total MEK1/2 and phosphorylated ERK1/2 (Thr202+Tyr204) over total ERK1/2 in the mESC lines harbouring the NF1$^{Phe1112Leu}$ and RAF1$^{Asp633Tyr}$ variants after EGF stimulation (100 ng/ml for 2 min.). (**B**) Significant reduction of phosphorylated MEK1/2 (Ser217/221) over total MEK1/2 and phosphorylated ERK1/2 (Thr202+Tyr204) over total ERK1/2 in the mESC lines harbouring the NF1$^{Phe1112Leu}$ and RAF1$^{Asp633Tyr}$ variants after insulin stimulation (100 nM for 10 min.). (**C**) Significant reduction of phosphorylated p70 S6K (Thr389) over total p70 S6K in the mESC lines harbouring the NF1$^{Phe1112Leu}$ and RAF1$^{Asp633Tyr}$ variants after insulin stimulation (100 nM for 10 min.). Error bars represent standard deviation. Data was analysed using a one-way ANOVA and Dunnett's post hoc test. *P < 0.05, **P < 0.01, ***P < 0.001, ****P < 0.0001.

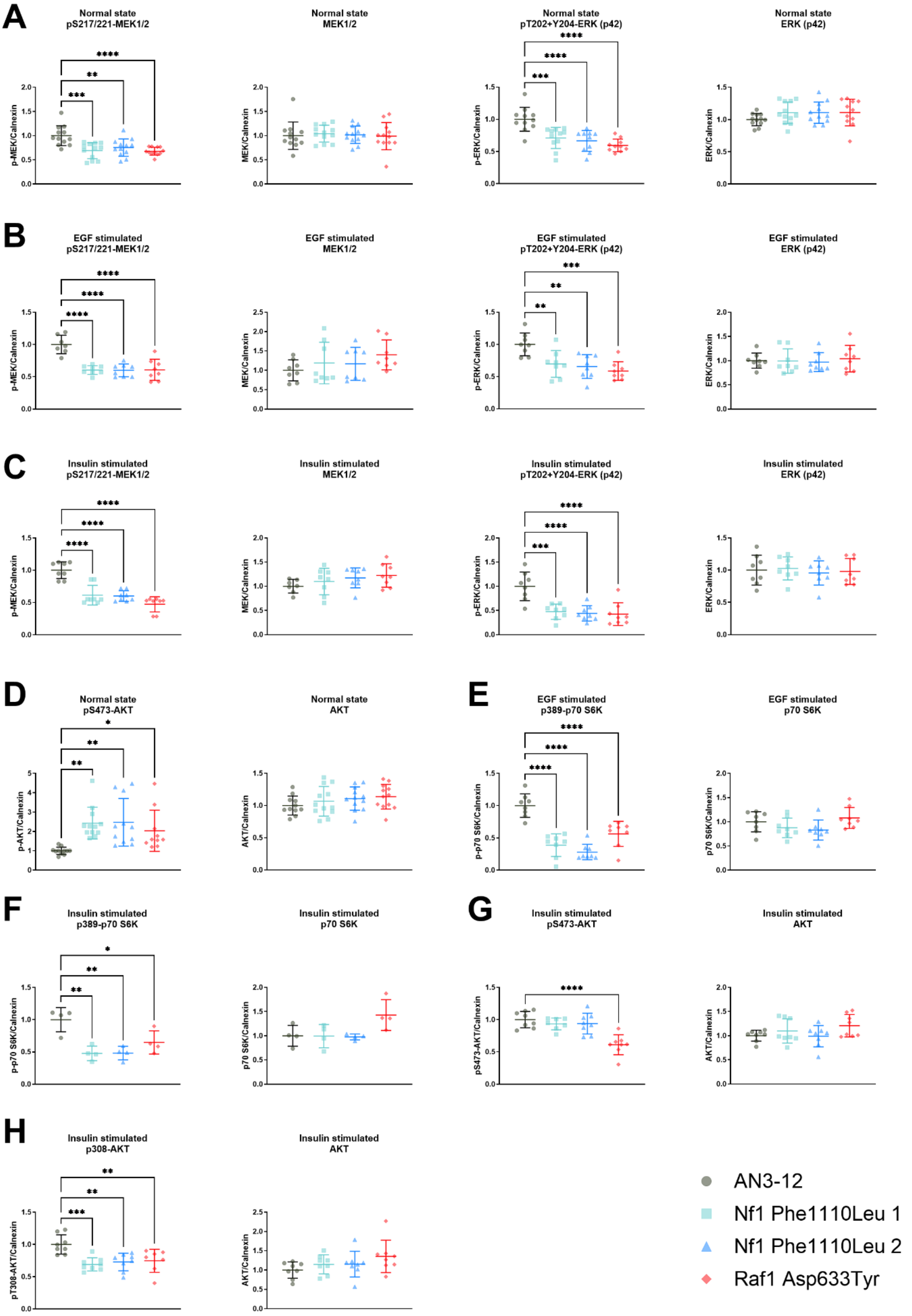


**Figure EV4 | Phosphorylated and total MAPK/ERK and IIS pathway-related protein levels in the NF1$^{Phe1112Leu}$ and RAF1$^{Asp633Tyr}$ variant mESC lines.**

(**A-C**) Significant reduction of phosphorylated MEK1/2 (Ser217/221) and phosphorylated ERK1/2 (Thr202+Tyr204), but not total MEK1/2 and ERK1/2, in the mESC lines harbouring the NF1$^{Phe1112Leu}$ and RAF1$^{Asp633Tyr}$ variants maintained in normal growth medium (**A**), as well as after EGF stimulation (100 ng/ml for

2 min.) (**B**) and insulin stimulation (100 nM for 10 min.) (**C**). (**D**) Significant increase of phosphorylated AKT (Ser473), but not total AKT, in the mESC lines harbouring the $NF1^{Phe1112Leu}$ and $RAF1^{Asp633Tyr}$ variants maintained in normal growth medium. (**E-F**) Significant reduction of phosphorylated p70 S6K (Thr389), but not total p70 S6K, in the mESC lines harbouring the $NF1^{Phe1112Leu}$ and $RAF1^{Asp633Tyr}$ variants after EGF stimulation (100 ng/ml for 2 min.) (**E**) and insulin stimulation (100 nM for 10 min.) (**F**). (**G**) Significant reduction of phosphorylated AKT (Ser473), but not total AKT, in the mESC line harbouring the $RAF1^{Asp633Tyr}$ variant. (**H**) Significant reduction of phosphorylated of AKT (Thr308), but not total AKT, in the mESC lines harbouring the $NF1^{Phe1112Leu}$ and $RAF1^{Asp633Tyr}$ variants after insulin stimulation (100 nM for 10 min.). Error bars represent standard deviation. Data was analysed using a one-way ANOVA and Dunnett's post hoc test. *P < 0.05, **P < 0.01, ***P < 0.001, ****P < 0.0001.

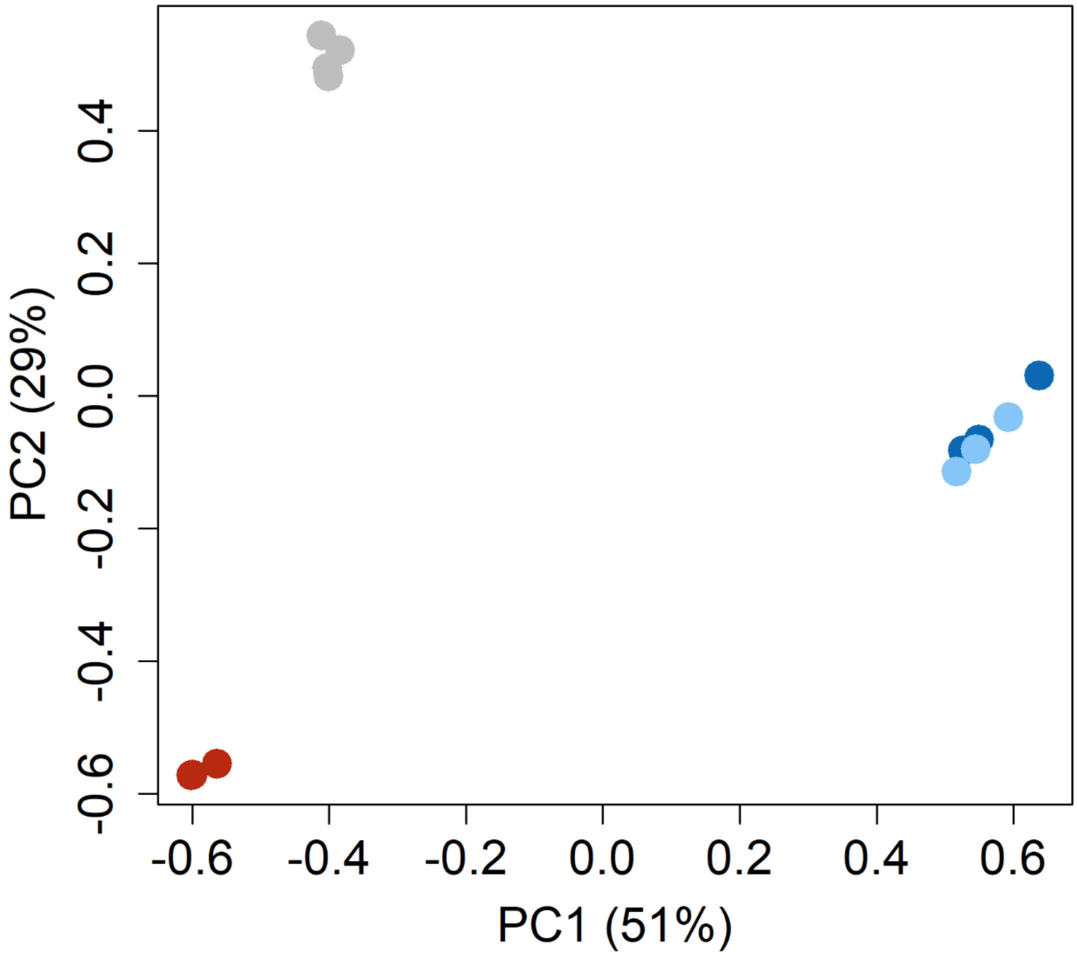


**Figure EV5 | Principal component analysis of proteomics data of the NF1$^{Phe1112Leu}$ and RAF1$^{Asp633Tyr}$ variant mESC lines.**

TMM normalised clustering confirms the two independent cell lines harbouring the NF1$^{Phe1112Leu}$ variant to be identical (light blue and dark blue dots). The RAF1$^{Asp633Tyr}$ cell line (red dots) clusters differently from the wildtype (grey dots) and NF1$^{Phe1112Leu}$ variant cell lines.

**Table EV1 | Gene list MAPK/ERK signalling pathway.**

| Ensembl stable ID | Gene name |
|---|---|
| ENSG00000078061 | *ARAF* |
| ENSG00000128272 | *ATF4* |
| ENSG00000157764 | *BRAF* |
| ENSG00000120129 | *DUSP1* |
| ENSG00000143507 | *DUSP10* |
| ENSG00000111266 | *DUSP16* |
| ENSG00000158050 | *DUSP2* |
| ENSG00000108861 | *DUSP3* |
| ENSG00000120875 | *DUSP4* |
| ENSG00000138166 | *DUSP5* |
| ENSG00000139318 | *DUSP6* |
| ENSG00000164086 | *DUSP7* |
| ENSG00000184545 | *DUSP8* |
| ENSG00000130829 | *DUSP9* |
| ENSG00000138798 | *EGF* |
| ENSG00000146648 | *EGFR* |
| ENSG00000134954 | *ETS1* |
| ENSG00000157557 | *ETS2* |
| ENSG00000139083 | *ETV6* |
| ENSG00000010030 | *ETV7* |
| ENSG00000177885 | *GRB2* |
| ENSG00000174775 | *HRAS* |
| ENSG00000133703 | *KRAS* |
| ENSG00000109270 | *LAMTOR3* |
| ENSG00000169032 | *MAP2K1* |
| ENSG00000126934 | *MAP2K2* |
| ENSG00000100030 | *MAPK1* |
| ENSG00000102882 | *MAPK3* |
| ENSG00000079277 | *MKNK1* |
| ENSG00000099875 | *MKNK2* |
| ENSG00000172680 | *MOS* |
| ENSG00000158186 | *MRAS* |
| ENSG00000196712 | *NF1* |
| ENSG00000213281 | *NRAS* |
| ENSG00000072062 | *PRKACA* |
| ENSG00000142875 | *PRKACB* |
| ENSG00000165059 | *PRKACG* |
| ENSG00000154229 | *PRKCA* |
| ENSG00000166501 | *PRKCB* |
| ENSG00000126583 | *PRKCG* |
| ENSG00000110786 | *PTPN5* |
| ENSG00000143851 | *PTPN7* |
| ENSG00000153233 | *PTPRR* |
| ENSG00000132155 | *RAF1* |
| ENSG00000116473 | *RAP1A* |
| ENSG00000127314 | *RAP1B* |
| ENSG00000109756 | *RAPGEF2* |
| ENSG00000145715 | *RASA1* |
| ENSG00000155903 | *RASA2* |
| ENSG00000058335 | *RASGRF1* |
| ENSG00000113319 | *RASGRF2* |
| ENSG00000172575 | *RASGRP1* |
| ENSG00000068831 | *RASGRP2* |
| ENSG00000152689 | *RASGRP3* |
| ENSG00000171777 | *RASGRP4* |
| ENSG00000117676 | *RPS6KA1* |
| ENSG00000071242 | *RPS6KA2* |
| ENSG00000177189 | *RPS6KA3* |
| ENSG00000162302 | *RPS6KA4* |
| ENSG00000072133 | *RPS6KA6* |

| **Ensembl stable ID** | **Gene name** |
| --- | --- |
| ENSG00000126458 | *RRAS* |
| ENSG00000262489 | *RRAS2* |
| ENSG00000160691 | *SHC1* |
| ENSG00000129946 | *SHC2* |
| ENSG00000148082 | *SHC3* |
| ENSG00000185634 | *SHC4* |
| ENSG00000115904 | *SOS1* |
| ENSG00000100485 | *SOS2* |

**Table EV2 | Results of the overrepresentation analysis using KEGG and MSigDB hallmark gene sets of the RAF1$^{Asp633Tyr}$ and NF1$^{Phe1112Leu}$ variant mESC lines.**

| Comparison | ID | Description | GeneRatio | BgRatio | p-value | FDR-adjusted p-value |
|---|---|---|---|---|---|---|
| RAF1$^{Asp633Tyr}$ Down | mmu01100 | Metabolic pathways | 294/1083 | 735/3366 | $2.45 \times 10^{-7}$ | $7.92 \times 10^{-5}$ |
| | mmu00982 | Drug metabolism - cytochrome P450 | 10/1083 | 11/3366 | $9.02 \times 10^{-5}$ | 0.015 |
| | mmu05323 | Rheumatoid arthritis | 13/1083 | 17/3366 | $2.21 \times 10^{-4}$ | 0.024 |
| | mmu04966 | Collecting duct acid secretion | 11/1083 | 14/3366 | $4.77 \times 10^{-4}$ | 0.039 |
| | mmu00480 | Glutathione metabolism | 19/1083 | 31/3366 | $7.65 \times 10^{-4}$ | 0.049 |
| | | HALLMARK_MTORC1_SIGNALING | 71/718 | 154/2209 | $1.83 \times 10^{-4}$ | 0.009 |
| | | HALLMARK_GLYCOLYSIS | 55/718 | 119/2209 | $9.32 \times 10^{-4}$ | 0.022 |
| RAF1$^{Asp633Tyr}$ Up | mmu04512 | ECM-receptor interaction | 20/842 | 29/3366 | $7.09 \times 10^{-7}$ | $2.18 \times 10^{-4}$ |
| | mmu03010 | Ribosome | 52/842 | 117/3366 | $2.47 \times 10^{-6}$ | $3.81 \times 10^{-4}$ |
| | mmu04514 | Cell adhesion molecules | 18/842 | 28/3366 | $1.19 \times 10^{-5}$ | 0.001 |
| | mmu05412 | Arrhythmogenic right ventricular cardiomyopathy | 19/842 | 32/3366 | $3.50 \times 10^{-5}$ | 0.002 |
| | | HALLMARK_MITOTIC_SPINDLE | 62/577 | 168/2209 | $8.88 \times 10^{-4}$ | 0.043 |
| NF1$^{Phe1112Leu}$ Down | mmu05323 | Rheumatoid arthritis | 14/863 | 17/3366 | $1.49 \times 10^{-6}$ | $4.77 \times 10^{-4}$ |
| | | HALLMARK_INTERFERON_GAMMA_RESPONSE | 32/600 | 69/2209 | $4.01 \times 10^{-4}$ | 0.019 |
| NF1$^{Phe1112Leu}$ Up | mmu05412 | Arrhythmogenic right ventricular cardiomyopathy | 20/915 | 32/3366 | $2.80 \times 10^{-5}$ | 0.009 |
| | mmu03010 | Ribosome | 51/915 | 117/3366 | $7.45 \times 10^{-5}$ | 0.012 |
| | mmu03008 | Ribosome biogenesis in eukaryotes | 33/915 | 69/3366 | $1.73 \times 10^{-4}$ | 0.018 |
| | | HALLMARK_MTORC1_SIGNALING | 68/649 | 154/2209 | $3.84 \times 10^{-5}$ | 0.001 |
| | | HALLMARK_MYC_TARGETS_V2 | 29/649 | 52/2209 | $5.22 \times 10^{-5}$ | 0.001 |

GeneRatio; size of the overlap of genes of interest with that node / size of the overlap of geneset with all members of the collection of geneset, BgRatio; genes of interest in the gene set / unique genes in the background genes.

**Table EV3 | Overview of used guide RNAs.**

| Target mutation | Vector backbone | gRNA1 | gRNA2 |
|---|---|---|---|
| NF1$^{Ala1127Val}$ | pSpCas9n(BB)-2A-GFP | GAAGATGAAAATGCACAAAC | CTTCACTACAGTCATTCAAA |
| NF1$^{Phe1112Leu}$ | pSpCas9n(BB)-2A-GFP | GAAGATGAAAATGCACAAAC | CATAAATAATGTGAAGTACC |
| RAF1$^{Asp633Tyr}$ | pSpCas9n(BB)-2A-GFP | GCACGCTGACTACATCCCCA | AGTGTGAGCTGCCCGATGCA |
| RRAS$^{Glu118Lys}$ | pSpCas9n(BB)-2A-GFP | TCTTGCAGTTTCAATGAGGT | GCATGAGGCACCACCTTAGC |

**Table EV4 | Overview of used single-stranded DNA oligonucleotides.**

| Target mutation | ssODN |
|---|---|
| NF1$^{\text{Ala1127Val}}$ | TCTCTGTATTTTCCAGGTACTTCACATTATTTATGAATCTTTTGAATGACTGTAGTGAAGTTGAAGACGAAAATG**T**TCAAACCGGTGGCAGGAAACGTGGCATGTCTCGGAGGCTGGCAT |
| NF1$^{\text{Phe1112Leu}}$ | CATGGTGTAAGGTCAGTCTCTGTATTTTACAGGTACTTCACATTA**C**TTATGAACCTTTTGAATGACTGTAGTGAAGTTGAAGACGAAAATGCTCAAACGGGTGGCAGGAAACGTGGCATG |
| RAF1$^{\text{Asp633Tyr}}$ | AAAATCAACAGGAGCGCCTCTGAGCCTAGCCTGCATCGGGCAGCTCACACTGAG**T**ACATCAATGCTTGCACGCTGACTACATCCCCAAGACTACCAGTCTTCTAGCTGATGATGTAGCTG |
| RRAS$^{\text{Glu118Lys}}$ | GGATGCTGGTGGGCCTGGCTGAGCACGGGGCAGGTTACTGCTAAGGTGGTACCTCATGCTACTGACTTCTCTTGCAAATTCAAT**A**AGGTGGGCAAGCTCTTCACACAGATCCTCAGAGTC |

The target mutations are highlighted in red, silent PAM and restriction site mutations are highlighted in orange.

**Table EV5 | Overview of oligonucleotides used for genotyping.**

| Target gene | Forward Primer | Reverse Primer |
|---|---|---|
| *Raf1* | TTAGATCATCTTCATGGTAGGCC | ACAGGGCCTTAACTTCATATTGC |
| *Nf1* | GTTGAAATTTTCCTTGTAATGAATC | TCCTTCCTCCTGGTTAATGG |
| *Rras* | GCAAGAGGAATTTGGTGCCA | GCAGGCAGGCAAGAAGTAAG |

**Table EV6 | Overview of primary antibodies used for western blotting.**

| Antigen | Source | Dilution | Vendor #ID |
|---|---|---|---|
| Phospho-p44/42 MAPK (Erk1/2) (Thr202+Tyr204) | Rabbit | 1:1000 | Cell Signaling #4370 |
| p44/42 MAPK (Erk1/2) total | Rabbit | 1:1000 | Cell Signaling #4695 |
| Phospho-Akt (Thr308) | Rabbit | 1:1000 | Cell Signaling #9275 |
| Phospho-Akt (Ser473) | Rabbit | 1:1000 | Cell Signaling #4060 |
| AKT total | Rabbit | 1:1000 | Cell Signaling #9272 |
| Phospho-p70 S6 Kinase (Thr389) | Rabbit | 1:1000 | Cell Signaling #9205 |
| p70 S6 Kinase total | Rabbit | 1:1000 | Cell Signaling #9202 |
| Phospho-MEK1/2 (Ser217/221) | Rabbit | 1:1000 | Cell Signaling #9154 |
| MEK1/2 total | Rabbit | 1:1000 | Cell Signaling #9122 |
| Phospho-CHK1 (Ser345) | Mouse | 1:1000 | Santa Cruz Biotechnology #8408 |
| CHK1 total | Rabbit | 1:1000 | Cell Signaling #2348 |
| Phospho-gamma-H2AX (Ser139) | Rabbit | 1:1000 | Abcam #11175 |
| gamma-H2AX total | Rabbit | 1:1000 | Cell Signaling #2577 |
| Calnexin | Rabbit | 1:1000 | Merck Millipore #208880 |

**Table EV7 | Overview of TaqMan Assays used for qPCRs.**

| Gene | TaqMan Assay ID |
|---|---|
| *Gapdh* | Mm99999915_g1 |
| *c-Myc* | Mm00487804_m1 |
| *Foxo3* | Mm01185722_m1 |
| *Nfe2l2* | Mm00477784_m1 |
| *Ets1* | Mm01175819_m1 |
| *Ets2* | Mm00468973_m1 |
| *Etv1* | Mm00514804_m1 |
| *Etv4* | Mm00476696_m1 |
| *Etv5* | Mm00465816_m1 |
| *Etv6* | Mm01261325_m1 |
